# Identification of pathogen genomic differences that impact human immune response and disease during *Cryptococcus neoformans* infection

**DOI:** 10.1101/592212

**Authors:** Aleeza C. Gerstein, Katrina M. Jackson, Tami R. McDonald, Yina Wang, Benjamin D. Lueck, Sara Bohjanen, Kyle D. Smith, Andrew Akampurira, David B. Meya, Chaoyang Xue, David R. Boulware, Kirsten Nielsen

**Author notes:** Corresponding author: Kirsten Nielsen, PhD, Department of Microbiology and Immunology, University of Minnesota, 689 23rd Ave SE, Minneapolis, MN 55455.

## Abstract

Patient outcomes during infection are due to a complex interplay between the quality of medical care, host immunity factors, and the infecting pathogen’s characteristics. To probe the influence of pathogen genotype on human immune response and disease, we examined *Cryptococcus neoformans* isolates collected during the Cryptococcal Optimal ART Timing (COAT) trial in Uganda. We measured human participants’ immunologic phenotypes, meningitis disease parameters, and survival. We compared this clinical data to whole genome sequences from 38 *C. neoformans* isolates of the most frequently observed sequence type (ST) ST93 in our Ugandan participant population, and an additional 18 strains from 9 other sequence types representing the known genetic diversity within the Ugandan *Cryptococcus* clinical isolates. We focused our analyses on 652 polymorphisms that: were variable among the ST93 genomes, were not in centromeres or extreme telomeres, and were predicted to have a fitness effect. Logistic regression and principal component analyses identified 40 candidate *Cryptococcus* genes and 3 hypothetical RNAs associated with human immunologic response or clinical parameters. We infected mice with 17 available KN99α gene deletion strains for these candidate genes and found that 35% (6/17) directly influenced murine survival. Four of the six gene deletions that impacted murine survival were novel. Such bedside-to-bench translational research provides important candidate genes for future studies on virulence-associated traits in human *Cryptococcus* infections.

**Author Summary:** Even with the best available care, mortality rates in cryptococcal meningitis range from 20-60%. Disease is often due to infection by the fungus Cryptococcus neoformans and involves a complex interaction between the human host and the fungal pathogen. Although previous studies have suggested genetic differences in the pathogen impact human disease, it has proven quite difficult to identify the specific C. neoformans genes that impact the outcome of the human infection. Here, we take advantage of a Ugandan patient cohort infected with closely related C. neoformans strains to examine to role of pathogen genetic variants on several human disease characteristics. Using a pathogen whole genome sequencing approach, we showed that 40 C. neoformans genes are associated with human disease. Surprisingly, many of these genes are specific to Cryptococcus and have unknown functions. We also show deletion of these genes alters disease in a mouse model of infection, confirming their role in disease. These findings are particularly important because they are the first to identify C. neoformans genes associated with human cryptococcal meningitis and lay the foundation for future studies that may lead to new treatment strategies aimed at reducing patient mortality.

## Introduction

*Cryptococcus neoformans* is the etiological agent of cryptococcal meningitis, the most common brain infection in Sub-Saharan Africa, which encompasses 15% of AIDS-related deaths [1]. As with all fungal pathogens, a major clinical concern is the small number of antifungal drug classes available (n=3) [2,3]. Researchers seek to identify the pathogen virulence factors that influence human health in order to develop novel drug targets to improve patient survival [4]. In addition to virulence factors that are common among all human pathogenic fungi, such as the ability to grow at 37°C, a number of *Cryptococcus-specific* virulence factors have been identified. The most well-studied include the polysaccharide capsule, the synthesis of melanin, and the secretion of extracellular enzymes such as phospholipases, laccase, and urease [5]. As we have previously discussed [6], there is not a clear quantitative association between *in vitro* virulence factor defects and clinical parameters of disease [7–13], thus studies clarifying this relationship are required.

Additional potential virulence targets have been identified through reverse genetic screens of the *C. neoformans* gene knockout collection [14]. A screen of 1201 knockout mutants from 1180 genes (20% of the protein coding genes) identified 164 mutants with reduced infectivity and 33 with increased infectivity in a screen for murine lung infectivity [7]. Deselarmos and colleagues [15] screened the same mutants for virulence in *Caenorhabditis elegans* and *Galleria mellonella* infection models and identified 12 mutants through a dual-species stepwise screening approach; all 12 also had attenuated virulence in a murine model (4 overlapped with those identified in the original murine lung screen). Many of the identified genes are associated with melanin production (which is not required for killing of *C. elegans*), thus the emerging picture is that genes that influence virulence are involved in multiple independent or parallel pathways such as melanization [15].

A complementary tactic to identify novel virulence factors is to use forward genetics, and look for association between strain background and virulence. *Cryptococcus* strains were originally classified by antigenic diversity, which led to differentiation into two species, *Cryptococcus neoformans* (var. *grubii* and var. *neoformans*, serotypes A and D, respectively) and *Cryptococcus gatti* (originally *C. bacillosporuus*, serotypes B and C [16]). The phylogenetic relatedness among strains has been subjected to a series of discussions that first used PCR fingerprinting and randomly amplified polymorphic DNA (RAPD) analysis [17] and then multilocus sequence typing (MLST) analysis [17] to classify strains based on sequence types (ST) defined in an online database (http://mlst.mycologylab.org). These analyses have led to competing species definition proposals. The first proposes classifying strains into seven species (two from *C. neoformans* following the serotypes and five from *C. gattii*) [18,19]. However, based on an analysis of 2600 strains, which revealed genetic diversity that is not-wholly captured by the seven species proposal [20] the second wants to maintain two groups, delineated as the “*C. neoformans* species complex” and the “*C. gattii* species complex” [21]. This “how do you define a species?” should not be written off as a purely philosophical issue [58], as we seek to discover whether there is a correlation between strain background and disease.

At a coarse level, there is a clear correlation between *Cryptococcus* variation and human infectivity. *C. neoformans* var. *grubii* strains cause the majority of infections in immunocompromised patients [22], while *C. gattii* is strongly implicated in cryptococcosis in immunocompetent individuals [23]. A handful of studies have demonstrated that there is also influence of phylogenetic relatedness on disease within var. *grubii* strains. The PCR/AFLP/MLST analyses divided var. *grubii* strains into three groups, VNI, VNII, and VNB strains. Beale and colleagues [10] found that among strains from South Africa, survival was lower for eight patients infected with VNB strains compared to the more common VNI or VNII strains (isolated from 175 and 47 patients, respectively). Similarly, Wiesner and colleagues [9] used MLST to type 111 strains isolated from Ugandan patients with their first episode of cryptococcal meningitis and conducted BURST clustering analysis to group strains with similar ST type (all of which are in the VN1 clade). BURST group 3 had significantly improved survival (62%) relative to BURST groups 1 and 2 (20% for both groups). Yet additional finer resolution studies by Mukaremera and colleagues within individual MLST sequence types (ST) show that there is also substantial variation in patient survival associated with individual strain differences [24]. Interestingly, while the South African clinical strains exhibited diversity in ST type, the Ugandan clinical strains were closely related, with ST93 strains accounting for approximately 60% of clinical isolates [9,10,24].

The overall picture that emerges from these studies is twofold. Strain background can significantly influence human disease, and there is tremendous disparity in strain frequency; some strain groups are much more common than others. ST93 is common in Uganda, but is also the most frequently isolated ST strain from HIV-infected patients in Brazil (85% [25,26]) and India (71% [27,28]). Sequence type prevalence also has a clear geographic component as different ST groups are dominant in other well-sampled countries (e.g., China, Thailand, Vietnam, Indonesia, Botswana, France [27–29]).

Here we sought to identify candidate genes associated with clinical phenotypes in human subjects. We took advantage of the large number of patients in Uganda infected with closely related ST93 strains and combined this with a powerful dataset collected during the Cryptococcal Optimal ART Timing (COAT) trial in Uganda [30]. When participants enrolled in the trial, strains were isolated and participant quantitative clinical and immunologic data were collected prior to treatment [40]. We sequenced the whole genomes of 38 ST93 strains, half from participants that survived the infection and half from participants that died, reasoning that restricting our search to variants among closely related strains would reduce background genetic noise. We conducted a series of statistical tests that identified 40 candidate genes and 3 hypothetical RNAs associated with clinical, immunologic, or *in vitro* phenotypes. We measured the virulence of 17 available KN99α knockout mutants for these genes in mice and found that 35% (6/17) had a significant association with mouse survival. Pathogen whole genome sequencing paired with statistical analyses of human clinical outcome data and *in vivo* virulence tests thus provides a new method to empirically probe the relationship between pathogen genotype and human clinical phenotype.

## Results

We whole-genome sequenced 56 *C. neoformans* VNI strains isolated from HIV-infected, ART-naive patients presenting with their first episode of cryptococcal meningitis at Mulago Hospital, Kampala, Uganda. The majority of strains (n=38) were chosen from ST93 isolates (the dominant genotype in Uganda [45]), collected as part of the Cryptococcal Optimal ART Timing (COAT) trial, where an array of human immunologic phenotypes and disease parameters were recorded for all participants. Approximately half of these strains were derived from participants who survived the infection (n=21) and half from participants who died (n=17). The remaining 18 strains were chosen to represent the diversity of the clinical strains in Uganda for phylogenetic purposes.

We identified 127344 SNPs and 15032 insertions/deletions (referred to as indels) associated with 7561 genes (or predicted genes) among the 56 sequenced *C. neoformans* strains. For ease of reference, we will refer to these SNPs, insertions, and deletions cumulatively as “variants”. Over three-quarters of the identified variants were non-coding variants not predicted to change the amino acid sequence of a gene: synonymous changes within the gene (22%), intergenic regions (3%), or designated as upstream or downstream of the associated gene (within 5kb of the nearest gene; 43% upstream, 10% downstream). The remaining (genic) variants are associated with 5812 different genes. Nonsynonymous coding changes are the largest class (90%) of these variants, with the remainder small insertion and deletion mutations.

The majority of genes have relatively few variants within the strain set, though 435 genes have over 50 variants (Figure 1A). There was a significant relationship between the number of variants and gene length (Pearson’s correlation test, t_4254_ = 33.001, *p*<0.001, cor = 0.45; Figure 1B), albeit with considerable variability around the line of best fit. The number of variants in each sequenced genome was extremely similar among strains from the same sequence type (Figure 1C), reflective of the phylogenetic distance from sequenced strains to the H99 reference genome (Figure 2).

**Figure 1.**
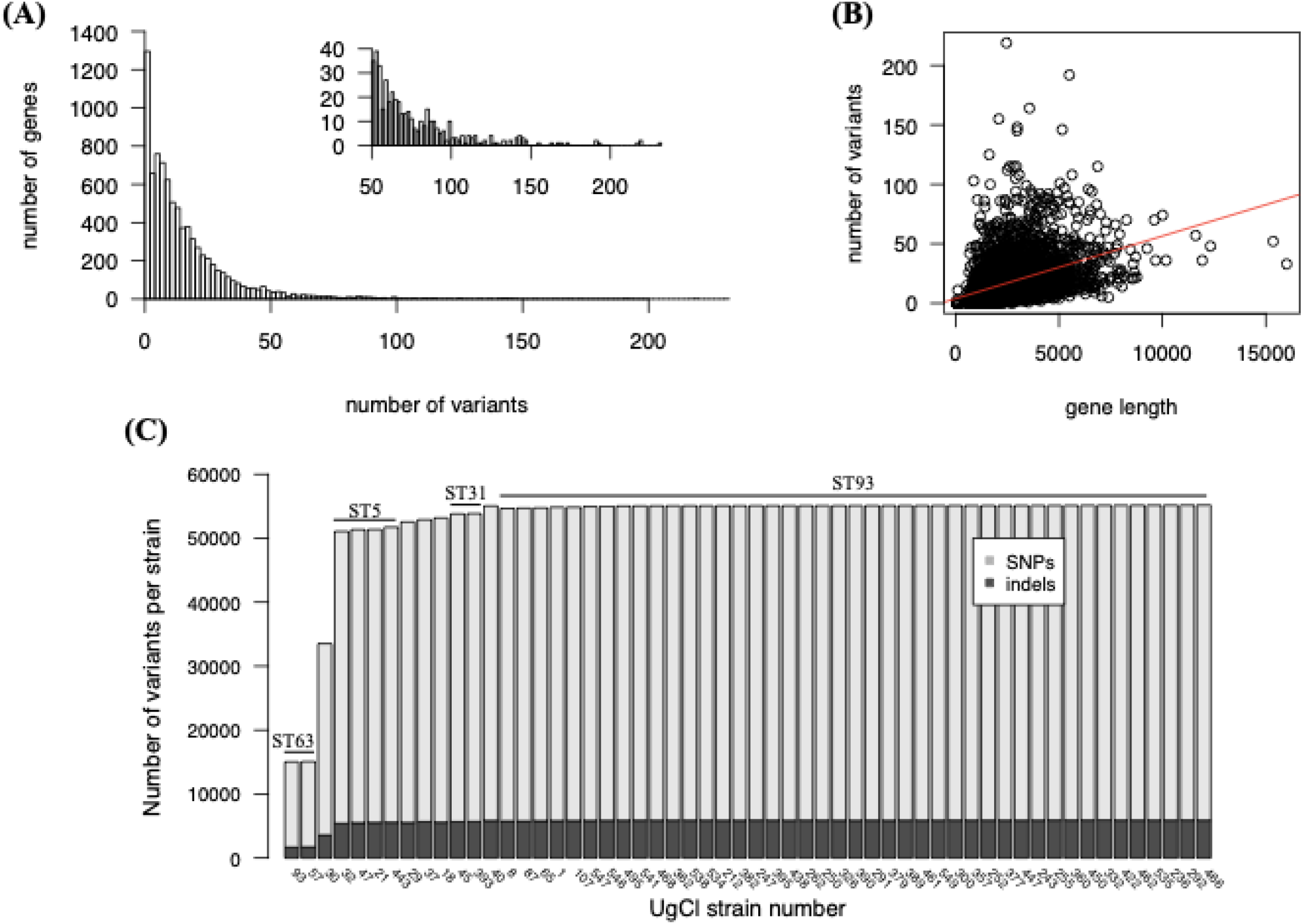
Variants identified among all strains. A) The number of variants per gene has a long right tail. The inset panel is the same data, zoomed for genes with at least 50 variants for visualization purposes. B) There is a significant and positive relationship between gene length and the number of variants per gene. C) The number of variants per strain matches the multi-locus sequence type (ST) among strains.

**Figure 2.**
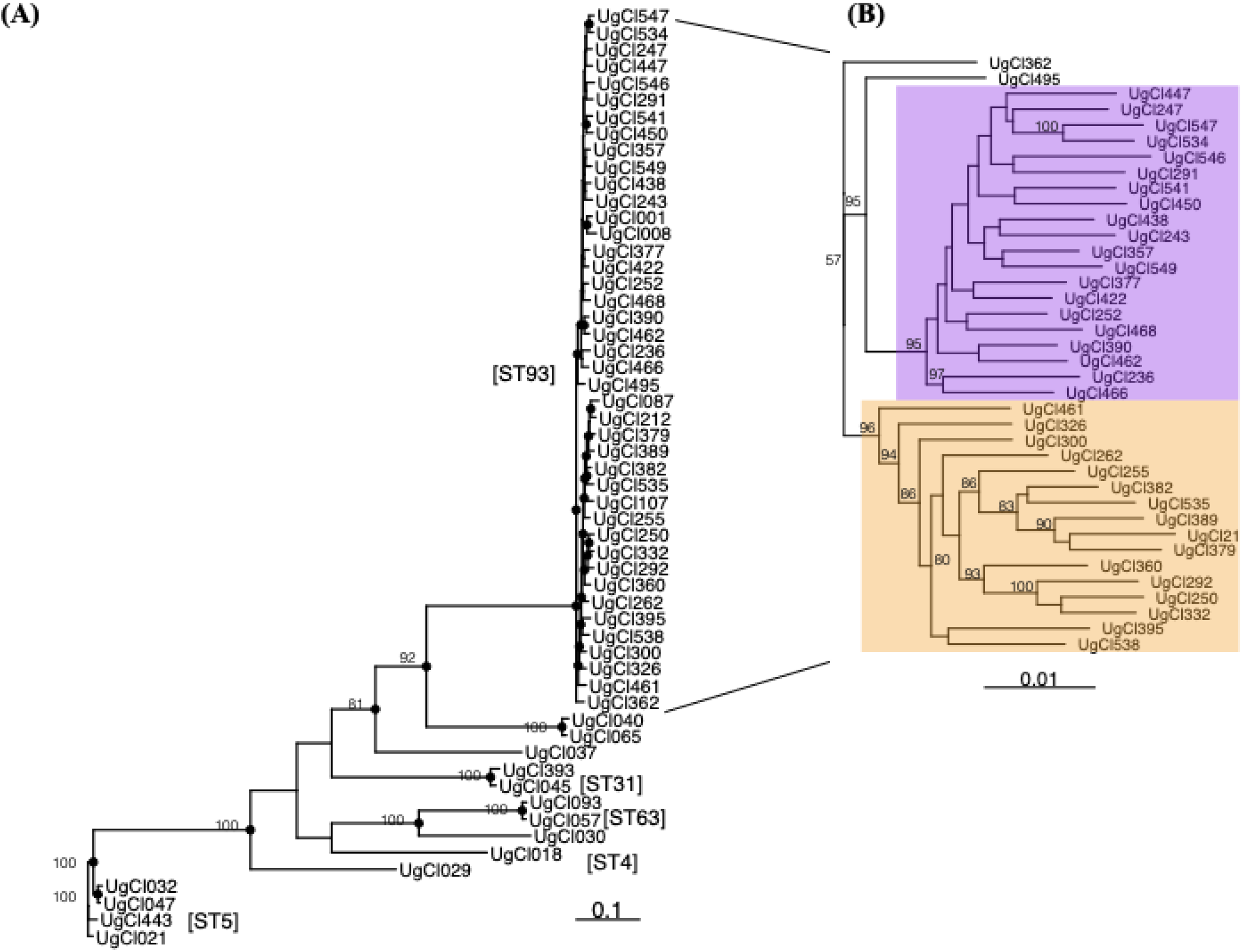
Phylogenetic analysis of all sequenced strains. A) The majority of ST93 strains fall into two well-supported clades, magnified in (B) for ease of viewing. ST93A (purple background) and ST93B (yellow background). Bootstrap values >50 are indiated with the numeric bootstrap value.

With this phylogenetic strain knowledge, we classified all variants into four categories: i) “common” variants differentiating Ugandan clinical isolates from the reference H99 genome; ii) “other” variants present only in non-ST93 genomes; iii) “allST93” variants present in all ST93 genomes but no other Ugandan ST genomes; iv) “someST93” variants present in some of the ST93 genomes. For our study, we considered the most interesting variants to be the “allST93” or “someST93” because these categories would potentially identify variants that could explain the increased overall pathogenesis of ST93 in humans (category iii), and will allow us to identify variants within ST93 associated with human clinical outcomes and phenotypes (category iv).

### Common variants in ST93

Variants that are in all ST93 strains and not the other sequenced strains (or the reference genome) can potentially tell us something about what differentiates strains in ST93 from other Ugandan strains. We identified 5110 variants common to all 38 ST93 genomes (4681 SNPs and 429 small indels). These variants were dispersed across the genome, associated with 2575 genes and 140 hypothetical RNAs (Figure 3, Table S1). The majority of these genes have one or a small number of variants, while a handful of genes had a very high number of variants (Table S2, 23 genes with at least 10 variants). The percentage of named genes in this set (8%, 2 of 24) matches the full gene set (8%, 686 out of 8338). The number of genes with a description (i.e., not “hypothetical protein” or “hypothetical RNA”) is actually lower in this gene set (33%) than the whole gene set (49%).

**Figure 3.**
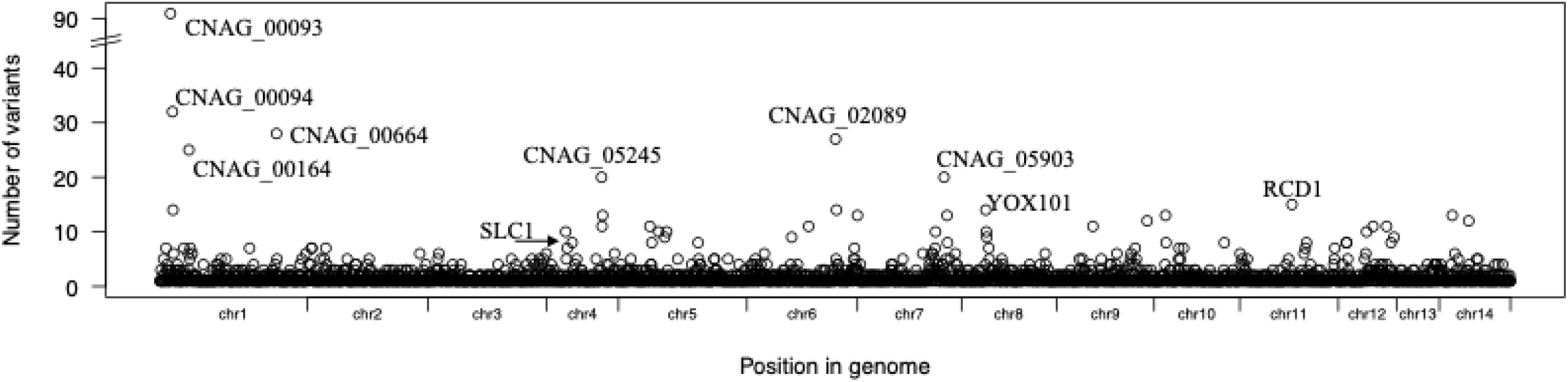
Variants that were common to all ST93 genomes are dispersed among 2715 genes and hypothetical RNAs. A small number of clustered genes have a large number of variants. In each cluster the gene with the highest number of variants is indicated. Genes with more than 20 variants and named genes are indicated. Table S2 lists all genes with 10 or more variants.

### ST93 clade-specific variants

Our primary aim was to identify the variants that are in some, but not all of the ST93 genomes, as these are the variants that can be used to examine genome associations with the measured human clinical phenotypes. When we examine the phylogenetic tree of ST93 COAT strains, we surprisingly identified a well-supported split between ST93 strains (Figure 2B), with 20 of the sequenced strains in one group (“clade A”), 16 strains in a second (“clade B”), and two ST93 strains outside of the primary clades. We identified 97 variants that differentiate strains in one clade from the other: 60 variants were unique to and in all clade A strains, and 37 variants were unique to and in all clade B strains. Clade-specific variants were located throughout the genome (Figure 4A) in 96 different genes. All except for one of the genes contained only a single clade-associated variant. In clade B, CNAG_06422 contains two variants in the 5’UTR that are three bases apart. An increased number of nonsynonymous and decreased downstream SNPs are observed in clade A compared to clade B (Figure 4B). Twenty-seven clade-specific mutations cause nonsynonymous amino acid changes (21 in clade A, 6 in clade B) and one small insertion mutation is present in clade A (Table S3). Although the majority of these variants are in genes that have not been characterized, four are in genes of known function: *LIV11* (CNAG_05422), a virulence protein of unknown function, *HSX1* (CNAG_03772), a high-affinity glucose transporter; *PTP2* (CNAG_05155), a protein tyrosine phosphatase; *SPT8* (CNAG_06597), and a predicted saga histone acetyltransferase complex component.

**Figure 4.**
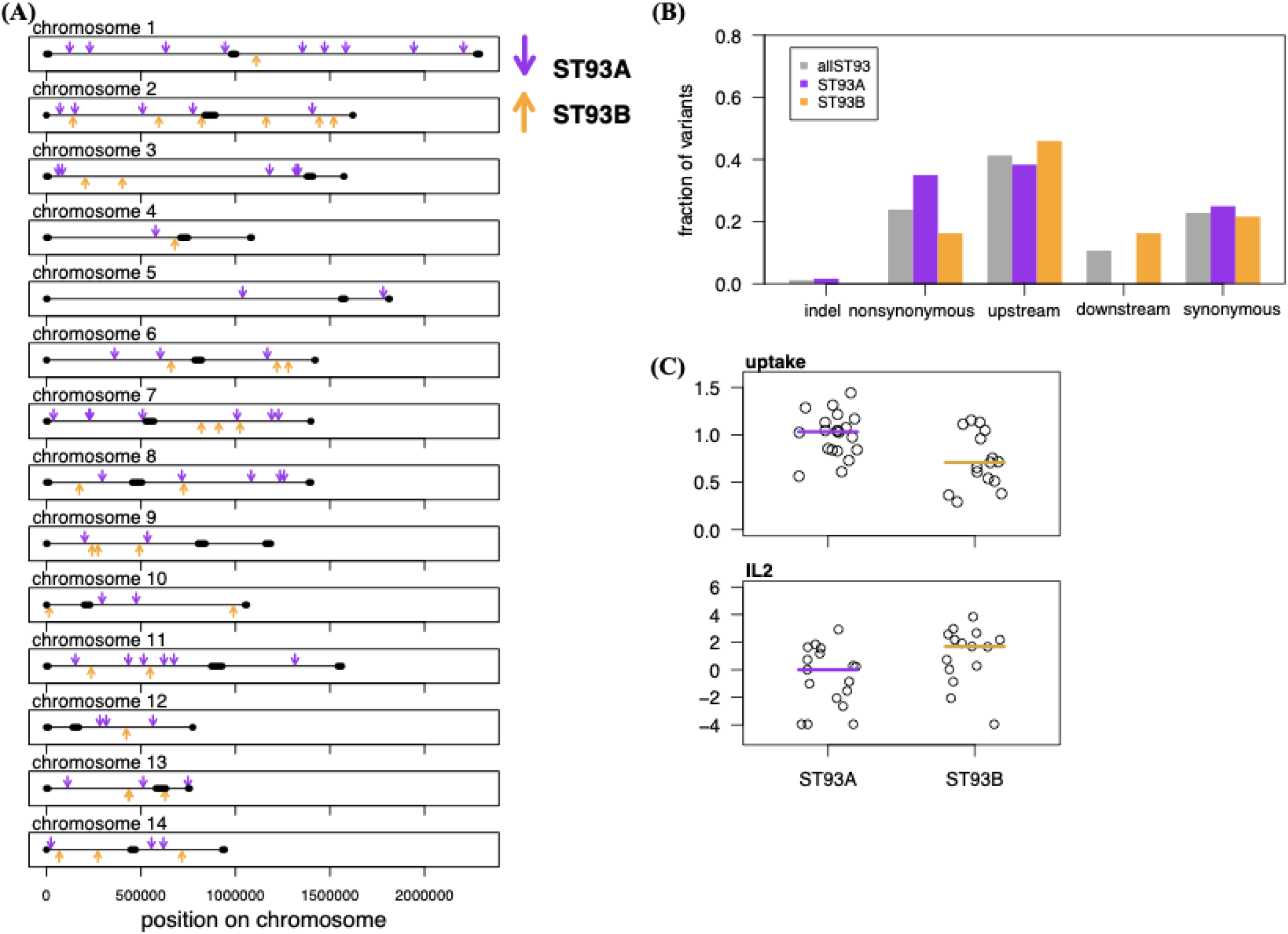
ST93A and ST93B clade-specific variants. A) Variants that are specific to the ST93A and ST93B clades are distributed across the genome. B) Upstream variants are the majority class found in all ST93 genomes (“ST93all”) and among the variants that are specific to either clade. By contrast, ST93A variants were more likely to be nonsynonymous and less likely to be downstream compared to ST93all or ST93B variants. C) IL2 cytokine levels in the CSF and *in vitro* macrophage uptake differed between ST93A and ST93B strains.

### Variant association with human clinical, immunologic and in vitro phenotypes

We next determined whether variants in the ST93 strains were associated with clinical measures of disease, CSF cytokines levels, or with *in vitro* phenotypes [30,40], (Table 1, see Methods for more details). We collectively refer to these three classes of phenotypes as “quantitative infection phenotypes”. We identified a significant correlation between the ST93 A/B clade with *in vitro* macrophage uptake rate and patient CSF interleukin (IL)-2 (non-parametric Wilcoxon rank sum test; uptake W = 226, *p* = 0.011; IL2 W = 66.5, *p* = 0.022; Figure 4C). There was not a significant relationship between ST93 clade and the other quantitative infection phenotypes (Figure S1A; non-significant t-test results in Table S4), nor between ST93 clade and survival (Figure S1B, Fisher-exact test, *p* = 0.33).

**Table 1.** Clinical Phenotypes measured from participants enrolled in the COAT trial and *in vitro* assays.

| Class | n | Phenotype Variable |
| --- | --- | --- |
| Clinical | 38 | CD4 T cell |
| Clinical | 35 | CSF white cell |
| Clinical | 31 | CSF protein |
| Clinical | 35 | HIV viral load |
| Clinical | 37 | CSF Clearance Rate (EFA) |
| Clinical | 30 | CSF CrAg LFA titer |
| Clinical | 38 | survival |
| Cytokines | 36 | IL1-β |
| Cytokines | 36 | IL-2 |
| Cytokines | 36 | IL-4 |
| Cytokines | 36 | IL-5 |
| Cytokines | 36 | IL-6 |
| Cytokines | 36 | IL-7 |
| Cytokines | 36 | IL-8 |
| Cytokines | 36 | IL-10 |
| Cytokines | 36 | IL-12 |
| Cytokines | 36 | IL-13 |
| Cytokines | 36 | IL-17 |
| Cytokines | 36 | G-CSF |
| Cytokines | 36 | GM-CSF |
| Cytokines | 36 | IFN-γ |
| Cytokines | 36 | MCP-1 |
| Cytokines | 36 | TNF-α |
| Cytokines | 36 | MIP-1β |
| *in vitro* | 38 | macrophage uptake |
| *in vitro* | 38 | macrophage adherence |
| *in vitro* | 37 | cell wall chitin |
| *in vitro* | 37 | absolute growth at 30°C |
| *in vitro* | 37 | fluconazole MIC |
| *in vitro* | 37 | amphotericin B MIC |

To examine associations between single variants and quantitative infection phenotypes (our primary objective), we parsed the 5605 variants that were in some (but not all) of the ST93 genomes. We took two complementary approaches to look for phenotypic associations. Our first tactic was to treat each measured phenotype as independent. For the second we used principal components analysis (PCA) to distill the 30 measured phenotypes into a smaller number of independent variables. Due to the nature of data collection for these types of phenotypic data, some strains were missing data for some phenotypes (Table S5). The most consequential was two strains missing data for all cytokine phenotypes.

For the first tactic we analyzed phenotypes in each class as independent datasets in a logistic regression approach (Figure 5). For each, we removed variants that were in very few (<4) strains, as well as those without a predicted function (i.e., synonymous and intergenic variants), and those that mapped to either the centromeric or extreme telomeric regions. This left us with 466 variants in 230 genes for the cytokine dataset and 652 variants in 328 genes for the clinical and *in vitro* datasets. For each dataset we then conducted logistic regression analyses for each variant against each phenotype and found that across all tests 207 variants from 115 different genes were significant for at least one phenotype. The majority (138 variants) were significant for a single phenotype. To partially correct for false positives, we focused our further analyses only on the variants that were significant for at least two phenotypes (“class **a**”) or when multiple significant variants were identified in the same gene (“class **b**”), or when the variant fulfilled both criteria (“class **ab**”). This narrowed the list to 145 variants from 40 genes and 3 hypothetical RNAs, with 13 variants in class **a**, 36 variants in class **b**, and 96 variants in class **ab** (Table 2, full information about significant variants including class in Table S6, full statistical information for each significant variant and phenotype in Table S7).

**Figure 5.**
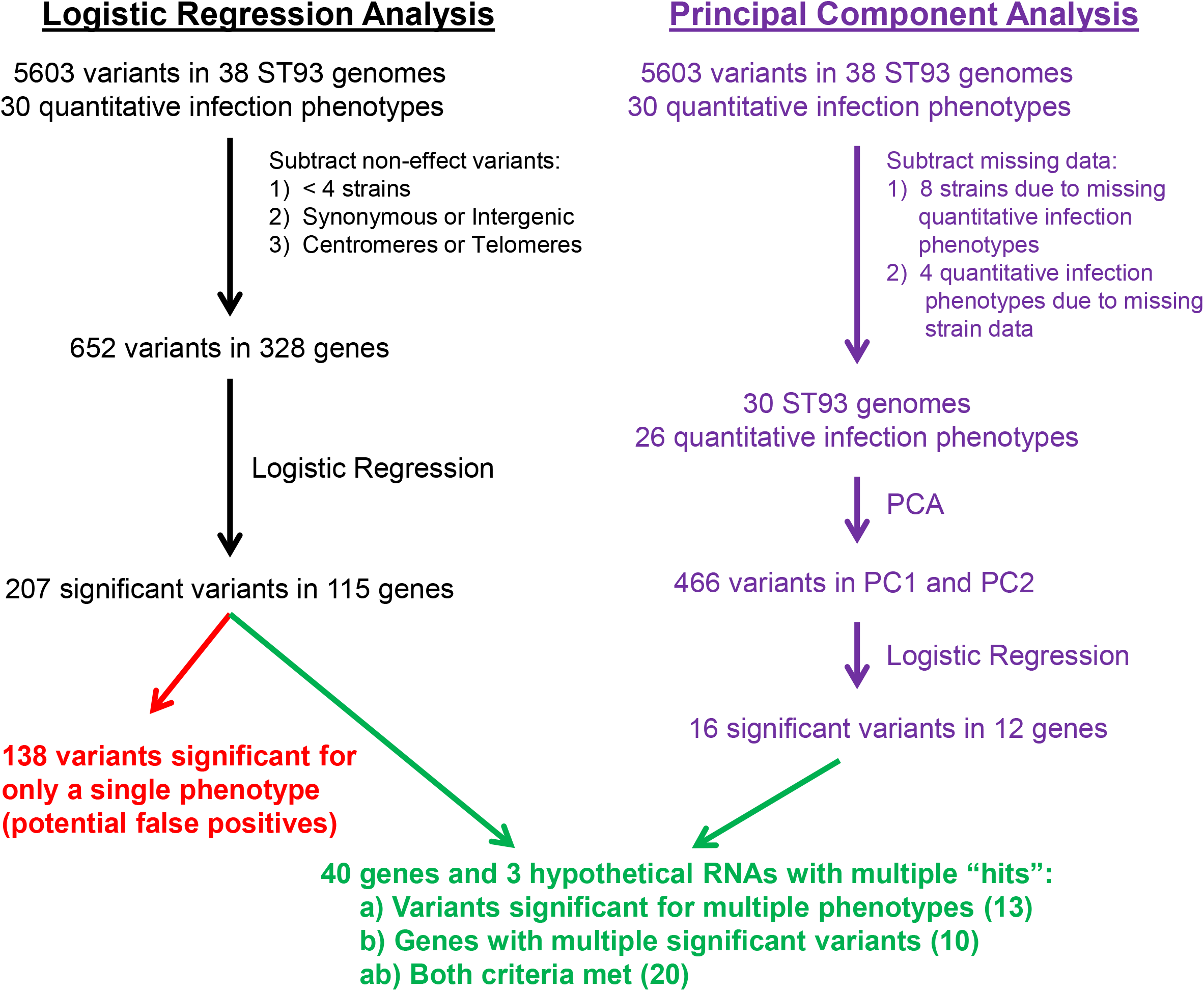
Flow chart for bioinformatic approaches used to identify *C. neoformans* genes associated with human infection. Two complementary approaches were used: logistic regression followed by cluster analysis and principal component analysis (PCA).

**Table 2.** Significant variants from linear regression analysis^a^.

| Gene<sup>b</sup> | Chr | Variant Positions | Effect<sup>c</sup> | Class<sup>d</sup> | Phenotypes |
| --- | --- | --- | --- | --- | --- |
| 00014 | 1 | 47564; 47575; 47671 | ns | b | GCSF; GCSF; GMCSF |
| 00363 | 1 | 927896; 927901 | ns | b | IL2; IL2 |
| 07950 | 1 | 975152; 975212;<br>975397 | up | ab | IL8, HIV<sub>1</sub>RNA; IL4, IL6, IL8, GMCSF, IFNγ, FLC; EFA |
| 06704 | 2 | 270700 | up | a | IL2, LFA |
| 02798 | 3 | 750294 | up | a | Protein, CD4, AMP |
| 05185 | 4 | 667433; 667446 | up | ab | Survival, uptake; uptake |
| 06876 | 5 | 7093 | down | a | IFNγ, MIP1β, TNFα |
| 01371 | 5 | 475470 | up | a | MCP1, HIV<sub>1</sub>RNA |
| 01241 | 5 | 836479; 836697;<br>836899 | up | ab | IL2; IL4, IL5, IL7, IL17, GMCSF, TNFα, chitin; IL5, IL12, IL13, IL17, GCSF, TNFα |
| 02475 | 6 | 221273; 221275;<br>221282 | up | ab | IL7, growth; growth; growth |
| 02176 | 6 | 988405; 988733;<br>988843; 989188;<br>989334; 989490;<br>989732; 990771;<br>990777; 990851;<br>990885; 991027 | down; ns;<br>ns; ns; ns;<br>ns; ns; ns;<br>ns; ns; ns;<br>up | ab | Chitin, SERT; IL1b, IL13, MCP1, MIP1β; MIP1β; IL12; AMP; HIV<sub>1</sub>RNA, SERT; IL2; IL10, MIP1β; MIP1β; IL10, MIP1β; SERT; IL13, TNFα, Survival |
| 02177 | 6 | 990701 | up | a | IL1β, IL6, IL10 |
| 02112 | 6 | 1160524; 1160528;<br>1160532 | up | b | AMP; AMP; AMP |
| 06525 | 7 | 11056; 14006 | ns ; up | ab | IL5, IL10; IL6,IL8 |
| 12610* | 7 | 49744 | up | a | MCP1, uptake |
| 06574 | 7 | 164473; 164887;<br>164926; 165027;<br>165704; 165873;<br>166309; 167135;<br>167224; 167292;<br>167370; 167687 | up | ab | HIV<sub>1</sub>RNA; IL2, TNFα; IL2, MIP1β; MIP1β; Survival, EFA; IL13; growth; IL13; GMCSF; IL1β, GCSF, MIP1β, uptake; CD4, uptake; protein |
| 05913 | 7 | 1205599; 1205600 | up | ab | MIP1β, adherence; IL13, IL17, MIP1β, adherence |
| 05937 | 7 | 1263610; 1263646;<br>1263647 | up | ab | Uptake, SERT; SERT; SERT |
| 07703 | 7 | 1341024 | ns | a | IL6, IL8 |
| 06968 | 8 | 1383765 | indel | a | IL12, IL17 |
| 04100 | 9 | 5213; 7729; 8171 | up | ab | adherence, FLC, SERT; growth; EFA, SERT |
| 04102 | 9 | 10033 | down | a | GMCSF, EFA |
| 04179 | 9 | 220963 | up | a | EFA, SERT |
| 04373 | 9 | 705343; 706175 | up | ab | IL8, EFA; survival |
| 04535 | 9 | 1115286 | up | a | IL17,GCSF,LFA |
| 07837 | 10 | 13558; 15288; 15302 | up; down;<br>down | b | IL2; WBCc; CD4 |
| 04922 | 10 | 18908; 18915; 18933;<br>18941; 18988; 18992;<br>18997 | up | b | IL2; IL2; IL2; IL2; adherence; adherence;<br>adherence |
| 08006 | 11 | 804710; 804742 | up | ab | IL4, IL5, IL6, MIP1β, TNFα, adherence,<br>chitin; IL4, IFNγ, MCP1, adherence |
| 01802 | 11 | 966644; 966669;<br>966700 | up | b | WBC; IL2; IL7 |
| 07026 | 12 | 11092; 11094; 11400;<br>11406; 11407; 11410;<br>11413 | up | ab | IL1β, IL13, survival, EFA; IL13, survival;<br>IL1β, IL7, IL13; IL1β, IL7, IL13; IL1β,<br>IL7, IL13; IL1β, IL7, IL13; IL1β |
| 05987 | 12 | 14009; 14035; 14125;<br>14197; 14202; 15014 | ns; ns;<br>indel; ns;<br>indel; up | ab | IL2; IL2; chitin; EFA, adherence; EFA,<br>adherence; adherence |
| 06169 | 12 | 502808; 502888;<br>502890; 503049;<br>503112; 503311;<br>503313; 503321;<br>503327; 503401 | down | ab | IL8; GMCSF, growth; IL6, IL8, GMCSF;<br>GMCSF, HIVrna; HIVrna, WBC; GCSF;<br>IL12, IL13, GCSF; IL12, IL13, GCSF,<br>MIP1β; IL12, IL13, MIP1β; IL10, chitin |
| 06256 | 13 | 11118; 11130 | up | ab; b | IFNγ, TNFα; TNFα |
| 13108* | 13 | 128625; 128715;<br>128729 | up | ab | IL13, GCSF; IL13, GCSF; IL13, GCSF |
| 06332 | 13 | 219021; 219311;<br>219312 | up | b | adherence; EFA; EFA |
| 06422 | 13 | 436551; 436554 | up | b | IL2; IL2 |
| 06490 | 13 | 655915 | indel | a | Protein, HIVrna, CD4 |
| 05450 | 14 | 342562 | ns | a | IL6, IL7, IL12, IL13, GCSF, MIP1β |
| 05661 | 14 | 908850; 908994;<br>909011; 909638;<br>910152; 910181 | up | ab | IL8, GMCSF, IFNγ, MCP1; uptake, FLC;<br>IL1β, IL8, MIP1β, uptake, FLC; adherence;<br>uptake; IL1β, IL6, IFNγ, HIVrna |
| 05663 | 14 | 910323; 910328;<br>910555 | down | ab | TNFα; IL1β, IL13, TNFα; survival |
| 05662 | 14 | 910742; 910822;<br>910834; 910926;<br>910939; 910964;<br>910966; 910979;<br>911099; 911129;<br>911206; 911262;<br>911292; 911308;<br>911321; 911352 | down | ab | AMP; survival, FLC; survival; SERT;<br>growth, SERT; survival, AMP; survival;<br>survival, uptake; IL12, GMCSF, growth,<br>TNFα, MCP1; IL12, IL13, IL17, MIP1β,<br>TNFα, growth, FLC, AMP, SERT; IL8,<br>MCP1, MIP1β; MCP1; IL2; adherence;<br>IL5; MCP1 |
| 13204* | 14 | 924025; 924047;<br>924049; 924050 | up | b | GMCSF; IL13; IL13; IL13 |

<sup>a</sup> Semicolons are used as separators between different variants. When only one effect is listed it is common among all variants in the gene.
- <sup>b</sup>Gene number corresponds to the CNAG number from the *Cryptococcus neoformans* H99 reference genome on FungiDB. Hypothetical RNAs are indicated with an \*. - <sup>c</sup>Effect designates location or type of variant: up, upstream of the coding region; down, downstream of the coding region; ns, nonsynonymous change in the coding region; indel, small insertion or deletion. - <sup>d</sup>Class type designations: a, genes with one variant significant for at least two phenotypes; b, multiple variants in the same gene with at least one significant phenotype each; ab, both criteria are fulfilled.

Following the default parameters in SnpEff, we used a very broad definition for calling variants upstream or downstream variants (+/- 5 kb). Over 80% of the significant variants were either upstream or downstream of genes (86 variants upstream, 34 variants downstream), with 20% within 1 kb (Table S6). Of the remaining variants, 21 were nonsynonymous, while 4 were indels. The majority of significant genes contained multiple significant variants (Table 2). In some cases, different variants in the same gene influenced the same phenotype, generally because the multiple significant variants are linked (e.g., three nonsynonymous variants in CNAG_00014 with the majority of ST93 strains falling into two haplotypes; one upstream SNP and two upstream insertions in CNAG_02112 with two haplotypes that influenced amphotericin B resistance). In other cases, such as CNAG_07950, there were six different haplotypes and three significant upstream variants that were associated with 8 unique phenotypes (IL8 was associated with the two variants, while HIVrna, IL4, IL6, GMCSF, IFNγ, Fluconazole MIC, and EFA were each associated with a single variant).

We also conducted PCA analysis as a second tactic to reduce the potential influence of phenotypic correlation on the results (Figure 5). As PCA requires complete datasets, we used data from the 27 phenotypes that had missing data from only three or fewer strains (i.e., we excluded CrAg LFA titer, HIV RNA viral load, CSF protein, and CSF white cell data, Table S5) and had to exclude 8 strains (UgCl212, UgCl332, UgCl357, UgCl422, UgCl447, UgCl461, UgCl541, UgCl549, Table 1). The ‘prcomp’ function from the R programming language was used to perform PCA on the two phenotypes which were scaled to have unit variance and shifted to be zero centered. We continued with the first two principal components by comparing the observed results to 20 datasets where the phenotypic data was randomized among strains (Figure S2A). Logistic regression analysis was run for each of the 466 variants that passed filter against PC1 and PC2. The PCA analysis yielded only 16 significant variants in 12 genes (Table 3). Only one of these genes, CNAG_07727, was not identified in the first analysis, and twelve of these variants were previously significant.

**Table 3.**
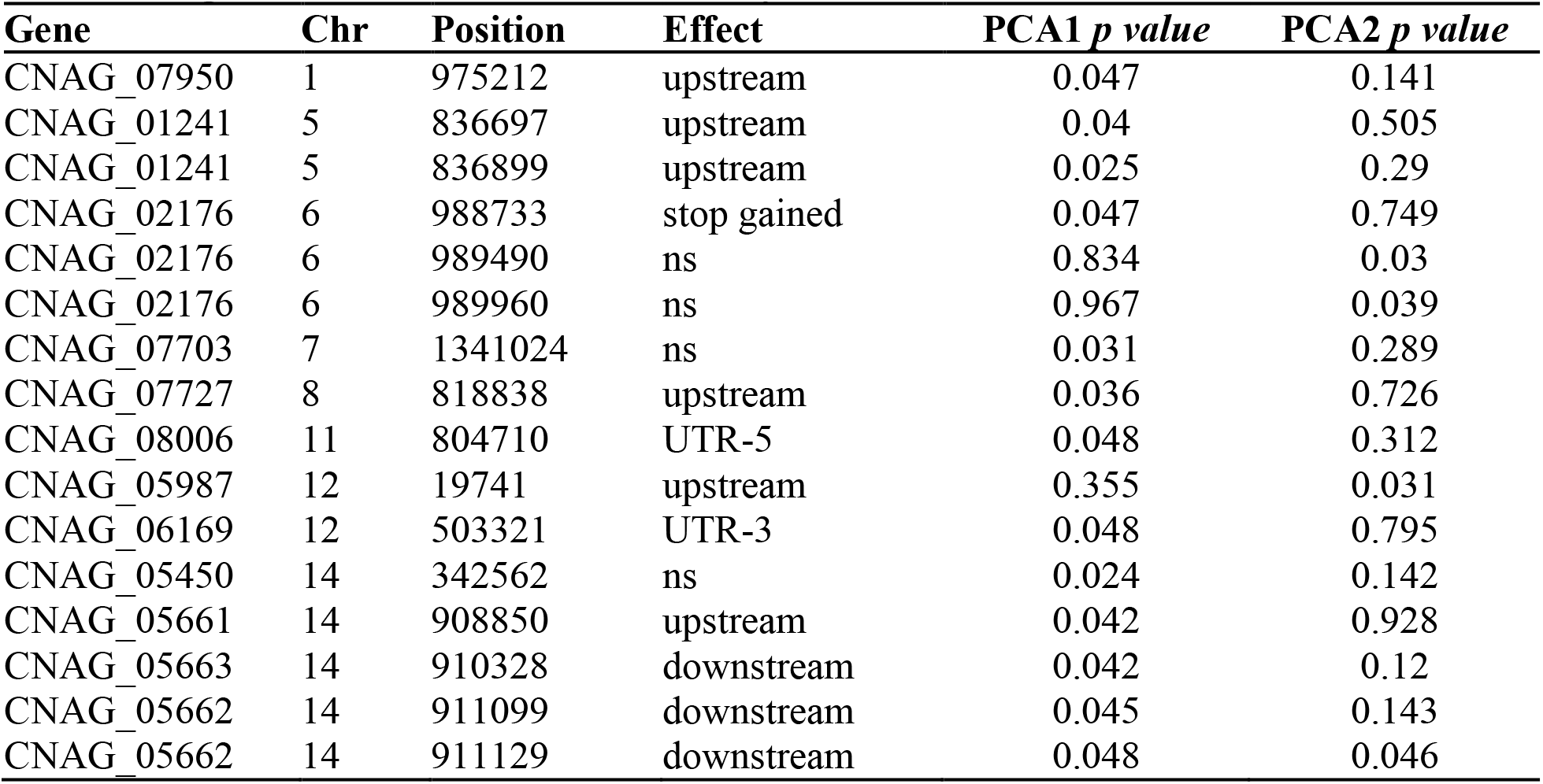
Significant variants from PCA analysis.

The majority of genes with a high number of significant variants were also genes with high numbers of sequenced variants and potentially-significant variants (Figure 6). In addition to variation among genes in regards to the number of significant variants within a gene (“sig variants”; ranging from 1-34), there was also variation in the number of variants that were identified within a strain (“sequenced variants”; range: 1-210) and the number of variants that passed our filters (“potentially-significant variants”: range: 1-32). This result highlights a limitation of genetic association screens such as the one we performed. Without additional biological validation it is difficult, if not impossible, to ascertain whether a given gene has many significant variants because of strong selection acting on that gene (e.g., if a knockout phenotype is beneficial there are many different positions that can reduce gene expression or protein levels) or because of relaxed selection and chance (i.e., if there is relaxed selection then many variants could be present, with statistical significance arising by chance). However, the fact that we do see areas of discordance between all the sequenced variants, potentially significant variants, and significant variants suggests many of our significant variants are not just a statistical artefact.

**Figure 6.**
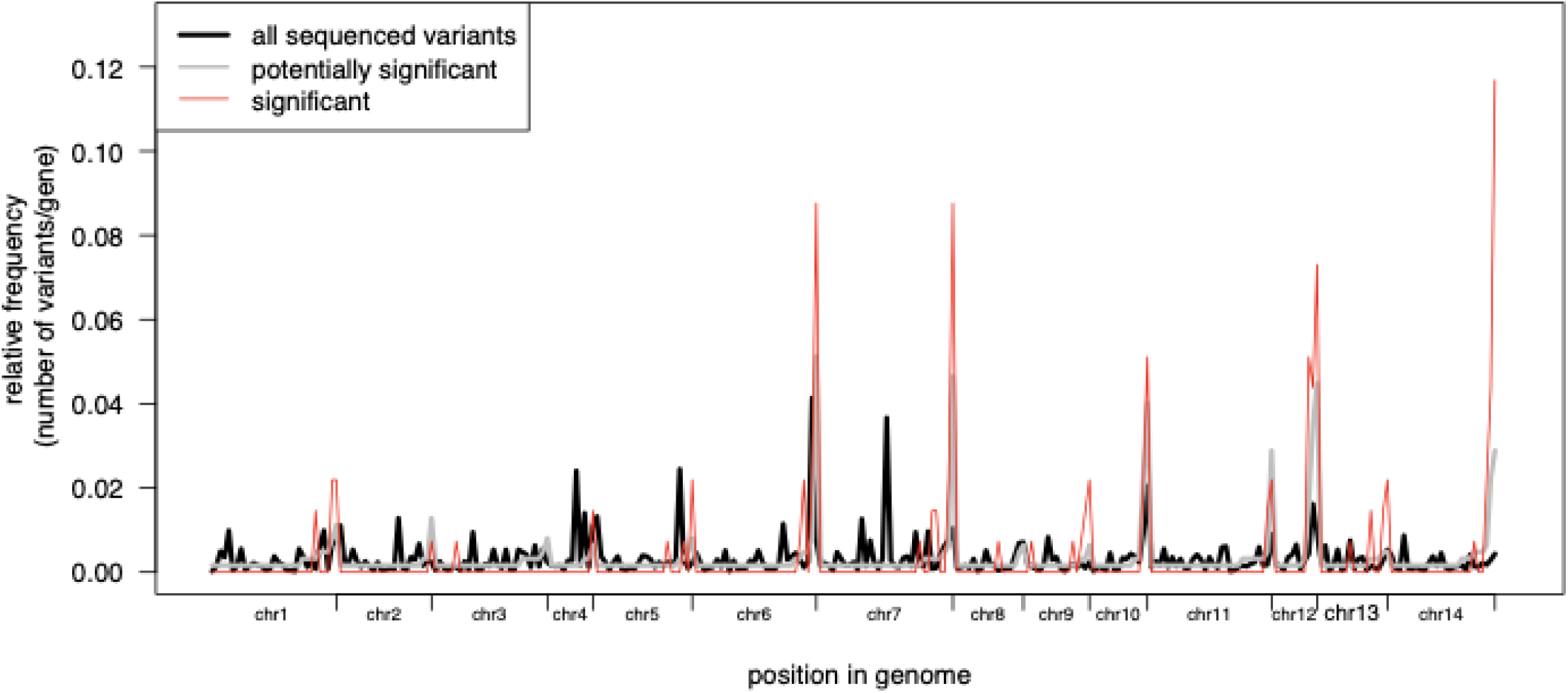
Comparison of variant frequency across the genome. The relative frequency of variants per gene for significant genes (red line) compared to all sequenced variants across all genomes (black line) and all variants within ST93 genomes (gray line). Only genes with at least one potentially significant variant are shown, hence the gray line does not reach 0. Discordance between the red compared to black and gray lines highlight areas with significant variants.

### In vivo virulence of identified genes

Our goal was to identify pathogen variants that impact human clinical disease phenotypes. Biological validation in humans is not possible. However, Mukaremera and colleagues recently showed that the mouse inhalation model of cryptococcosis accurately recapitulates human infections and can be used to dissect *C. neoformans* genetic factors that influence human disease. [24]. Thus, as a first step to probe the biological significance of the genes identified in our analyses, we tested the virulence of 17 available KN99α deletion strains in the inhalation mouse cryptococcosis model. Six (35%) of the tested deletion strains had a significant virulence effect on mouse survival compared to the control KN99α strain: three strains had increased virulence (CNAG_02176, CNAG_06574, CNAG_06332) and three strains had decreased virulence (CNAG_06986, CNAG_04922, CNAG_05662) (statistical results in Table 4, significantly different strains in Figure 7, non-significant strains in Figure S3). Although gene deletion mutants are only one way to biologically probe whether a candidate gene has a true virulence phenotype, we did find that the number of significant variants in a gene (Table 2) was a significant predictor of the deletion mutations having a virulence effect (linear model, F1, 15 = 8.493, *p* = 0.011).

**Table 4.**
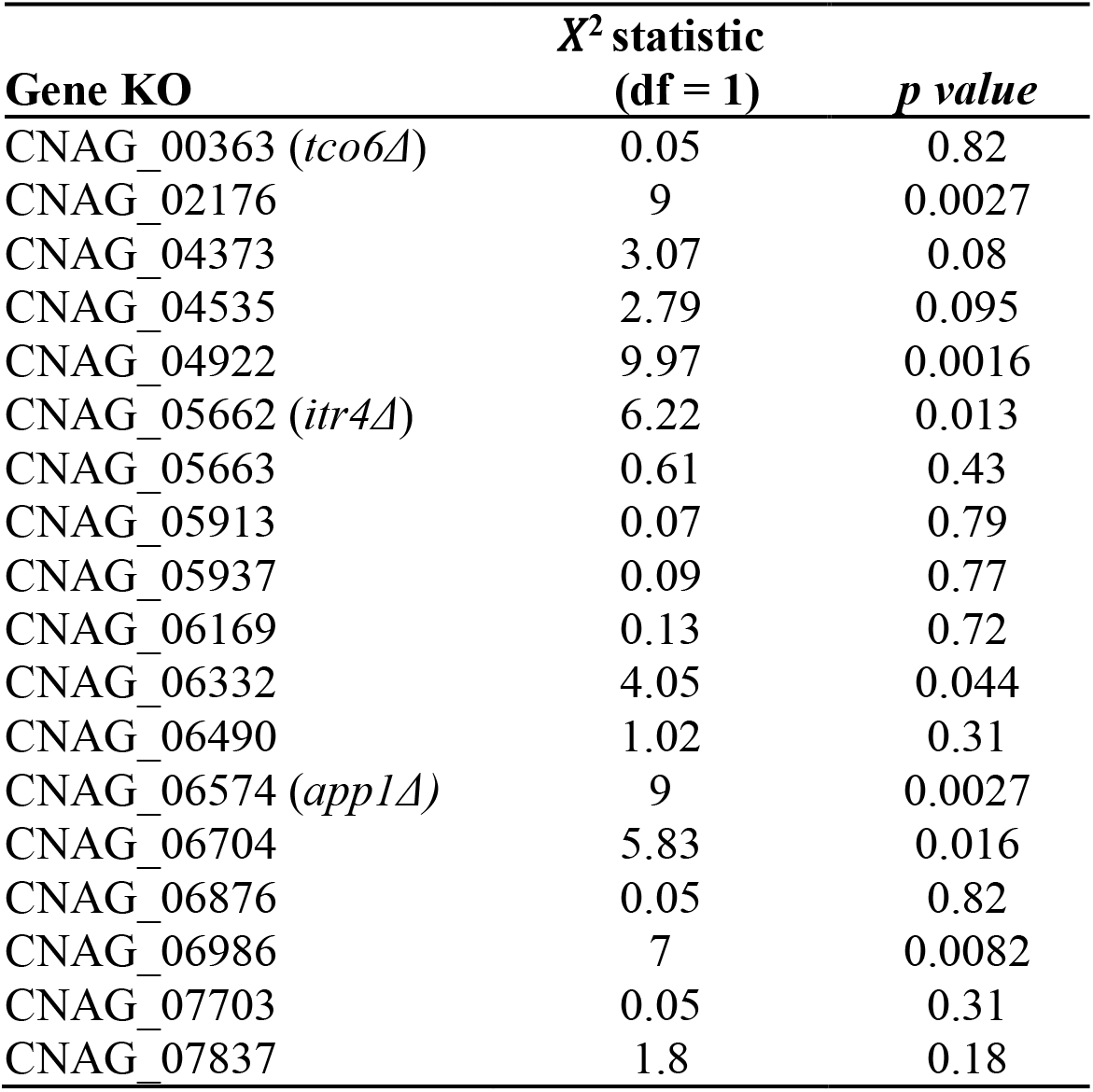
Survival curve statistical results.

**Figure 7.**
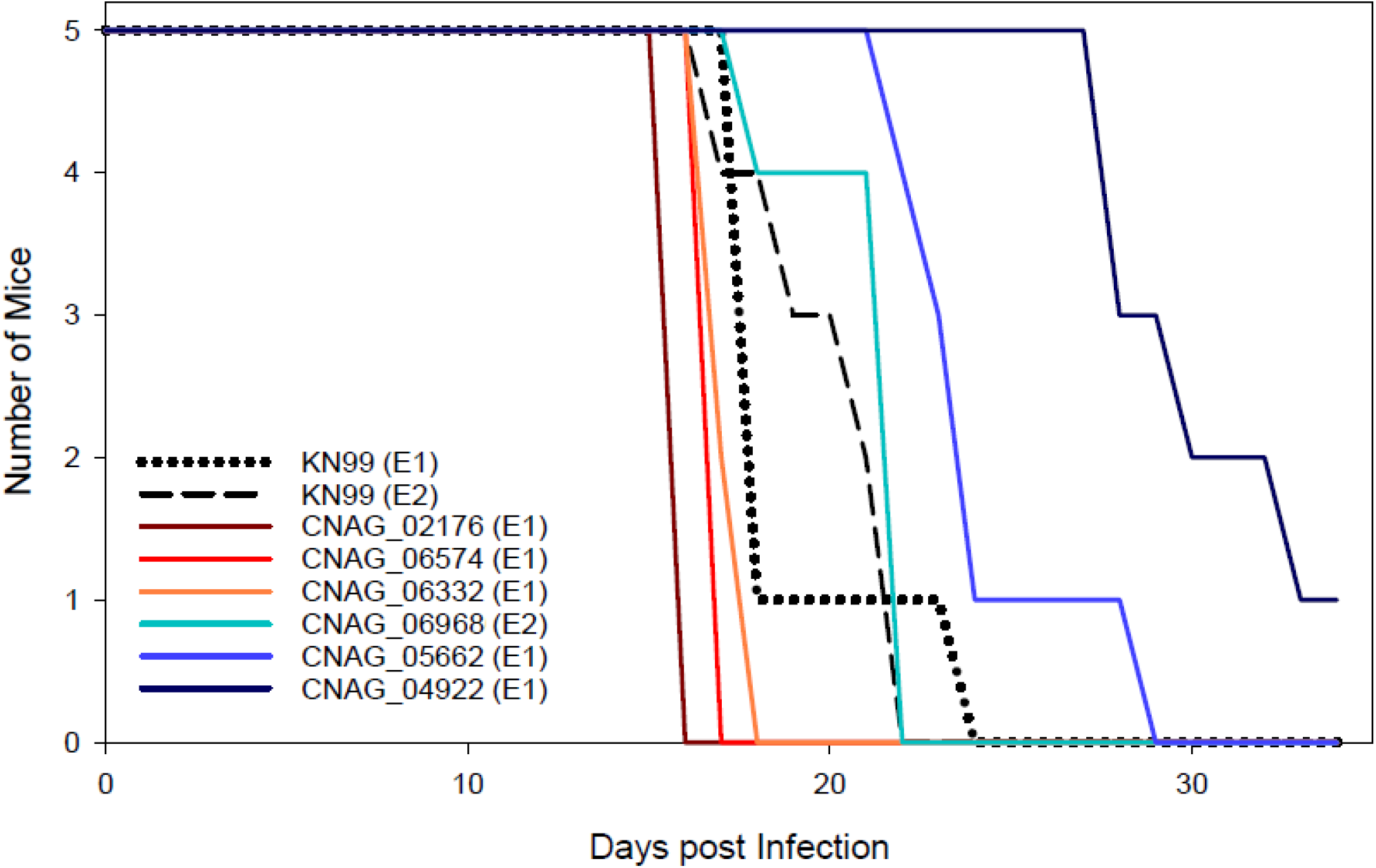
Deletion strain virulence in mice. Groups of five 6-8 week old C57Bl/6 mice were infected intranasally with 5 × 10^4^ cells. Progression to severe morbidity was monitored for 35 days and mice were sacrificed when endpoint criteria were reached. Strains were tested in two separate experiments indicated as E1 or E2, respectively.

### In vivo and in vitro analysis of itr4Δ and clinical strains

The gene with the highest number of significant variants in our candidate gene list was CNAG_05662 (*ITR4*), which has been reported as a member of the inositol transporter gene family [59]. The *itr4Δ* mutant strain had reduced virulence in the mouse model whereas the *itr4Δ*:ITR4 complement strain had equivalent virulence to the laboratory reference background strain KN99α showing that the ITR4 deletion is responsible for the virulence defect in the *itr4Δ*mutant (Figure 8A). In this lower inoculum experiment, three of the *itr4Δ* infected mice survived until the experiment was ended on day 44 (Figure 8A). Terminal colony forming units (CFUs) from the brain and lungs of the survivors showed complete fungal clearance in one mouse and a low fungal burden in the lungs (2 × 10^2^ CFUs) in the second mouse. The third mouse had 5.64 × 10^5^ CFUs in the lungs and 1.35 × 10^4^ CFUs in the brain. Evaluation of the fungal burden at seven days post-infection showed more *itr4Δ* mutant CFUs in the lungs than KN99*α* and *itr4Δ:ITR4*, and no *itr4Δ* CFUs in the brain (Figure S4), suggesting the reduced pathogenesis observed in the *itr4Δ* mutant is likely due to reduced growth in or delayed dissemination to the brain.

**Figure 8:**
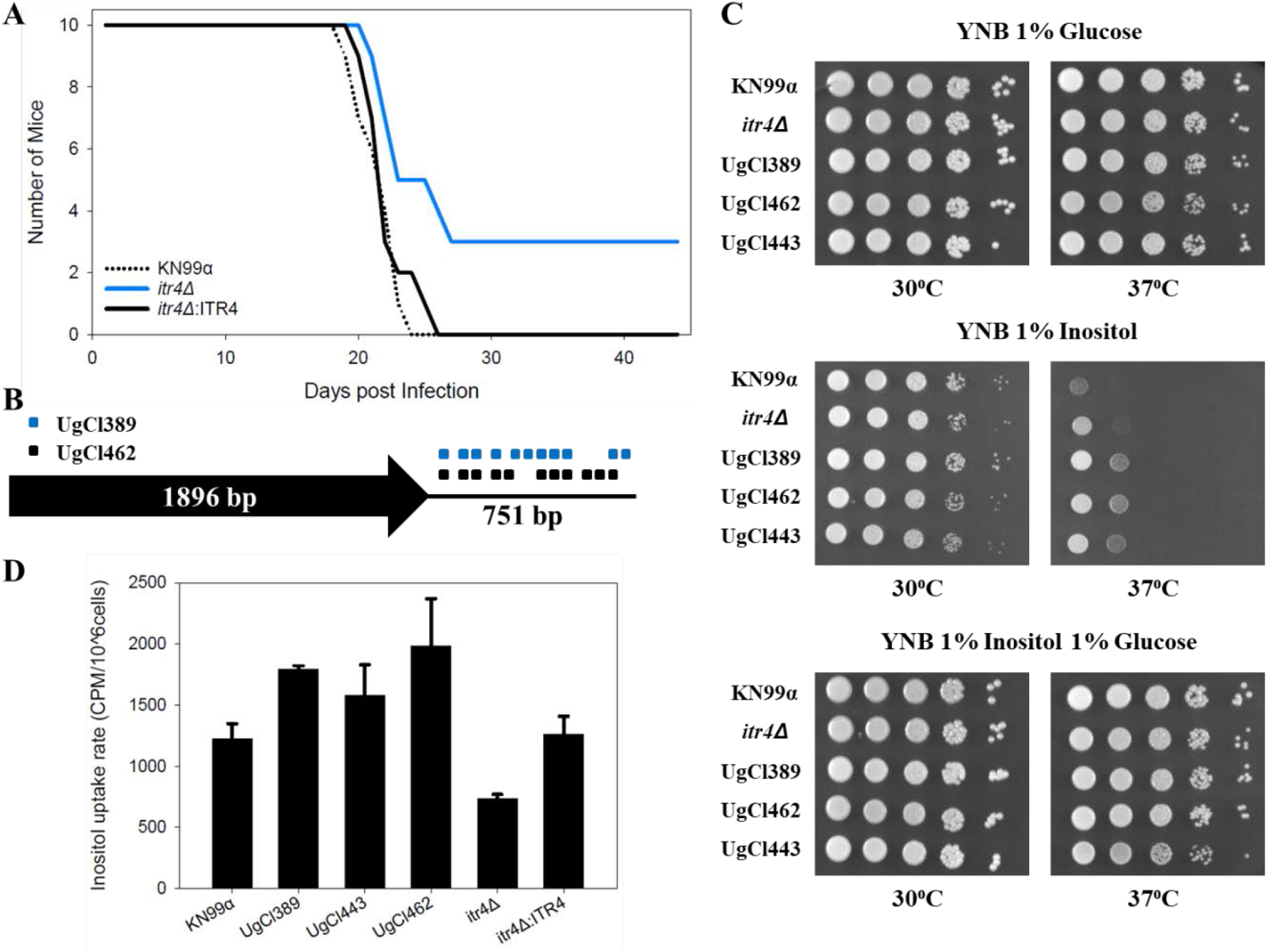
Analysis of *ITR4* through *in vivo* virulence and *in vitro* growth and inositol uptake. A) Groups of ten 6-8 week old C57Bl/6 mice were infected intranasally with 1 × 10^3^ cells. Progression to severe morbidity was monitored for 44 days and mice were sacrificed when endpoint criteria were reached. B) Schematic diagram showing location of the variants in the UgCl389 and UgCl462 clinical isolates relative to the ITR4 coding region. UgCl443 has the H99 reference allele. C) Growth assay of *C. neoformans* wild type strain *KN99α, itr4Δ* mutant, and clinical strains on medium with different inositol levels. Yeast cells were cultured in YPD medium. Equal cell concentrations were spotted as 10-fold serial dilutions onto YNB plates made with 1% glucose, 1% inositol, or 1% glucose and 1% inositol. Plates were incubated at 30°C and growth was examined after 4 days. The assay was repeated three times with similar results. D) Inositol uptake analysis of *C. neoformans* strains. Yeast cells were mixed with 3H-labeled inositol and incubated at 30°C for 10 minutes in triplicate and repeated twice with similar patterns. Error bar indicates the standard deviation of the three replicates.

To further determine the role of the genetic variants in the biological function of *ITR4*-KN99*α*, *itr4Δ*, and three clinical strains (UgCl389, UgCl462, and UgCl443) were tested for growth with inositol and inositol uptake. The variants associated with the *ITR4* locus in these clinical strains are proximal to the coding region – both UgCl389 and UgCl462 have 11 single nucleotide polymorphisms immediately downstream of the coding region whereas UgCl443 contains the H99 reference allele for *ITR4* (Figure 8B). All the clinical strains showed enhanced growth with inositol compared to KN99*α*, and similar to *itr4Δ* (Figure 8C). UgCl389 and UgCl462 were also more efficient at inositol uptake, while UgCl443 was similar to KN99*α*, and *itr4Δ* had decreased inositol uptake (Figure 8D). Taken together, these data highlight the complex nature of the multiple variants across the clinical strains. Due to differences in genetic background between the clinical strains, interpretation of the impact of specific variants and/or gene alleles is challenging.

## Discussion

Virulence is a multifaceted phenotype, as many different pathogen and host characteristics will determine the severity of a given infection. Here we paired a powerful dataset from the Cryptococcal Optimal ART Timing (COAT) trial in Uganda with pathogen whole genome sequencing technology to identify candidate *C. neoformans* genes that were statistically associated with quantitative human infection phenotypes. The technique of using Genome-wide Association Studies (GWAS) to uncover genic variants linked to disease was developed fourteen years ago in the context of human disease genetics [46]. Here we looked for association between variants within 38 ST93 *C. neoformans* isolates from participants enrolled in the COAT trial with 30 measured clinical phenotypes, cytokines, and *in vitro* phenotypes. We took two complementary tactics to identify candidate genes. The first treated each measured phenotype as independent, yet only included genes that either have a variant significantly associated with multiple phenotypes (13 genes), genes with multiple significant variants (10 genes), or both (20 genes). We also conducted a PCA analysis examining the first two principal components from a PCA on the 27 phenotypes and 30 strains with sufficient data. The resultant reduction of power is unfortunate, but not surprising when dealing with human data, and the detrimental impacts of missing clinical data have been previously discussed [47], and indeed is why we took both tactics. The PCA analysis yielded a total of 12 genes, 11 that overlapped with those identified in the first analysis and one additional gene. Combined, we identified 40 candidate *C. neoformans* genes and three hypothetical RNAs associated with infection phenotypes among the ST93 strains.

The statistical analysis is blind to any prior knowledge of the genes, and thus does not depend on prior annotation. Accordingly, the majority of genes we identified have not yet been named and roughly half (19) are listed as “hypothetical proteins” on FungiDB. Interestingly, only two of these 19 genes are conserved among fungal taxa, and curating information about orthologues from FungiDB (https://fungidb.org/fungidb/) suggests that the majority of others are either unique to *C. neoformans* or only have orthologues in the very closely related species complex *C. gattii* (Table S8). This is consistent with the logic of Liu et al. [7], who purposefully targeted genes that did not have homologues in *Saccharomyces cerevisiae* during the construction of the original H99 gene deletion collection [14] (an 1180 gene collection in *C. neoformans* H99, which corresponds to ~20% of the protein coding genes).

We took advantage of the newer KN99*α* gene deletion collection [48] and found that 35% (6/17) of the available gene knockouts had an effect on virulence in mice. The significant genes with a virulence change in mice include two named genes *ITR4* (CNAG_05662) and *APP1*(CNAG_06574) as well as three hypothetical proteins from closely related species (CNAG_02176, CNAG_04922, CNAG_06332) and one hypothetical protein with broad taxonomic distribution (CNAG_06968). *APP1* is a cytoplasmic protein involved in extracellular secretion and reduced phagocytosis. The *app1*Δ mutant has previously been shown to have decreased virulence in mice [49]. Interestingly, this is opposite from our mouse model that showed increased virulence of the *app1*Δ mutant. This difference could be due to the differential immune response in BALB/C (previous study, type 1 immune response) and C57Bl/6 mice (current study, type 2 immune response) that likely gives a hint to the mechanism of *APP1* in human disease.

Intriguingly, *ITR4* (synonym *PTP1*) was the top hit in a screen that identified genes that were overexpressed in an intracellular environment (amoebae and murine macrophages) compared to the lab medium YPD [50]. In that study, *itr4*Δ did not differ from wildtype in mice or *Galleria mellonella* virulence assays [50], though these studies were performed in a different genetic background from our KN99*α* reference strain. Using gene complementation, we clearly show the virulence defect in the *itr4*Δ mutant is due to deletion of the *ITR4* gene. However, our analysis of differences in growth and uptake of inositol in clinical isolates with different variants was less conclusive. All of the clinical strains appeared to be better adapted for growth and uptake of inositol compared to the KN99*α* reference strain. This is not surprising, given that the clinical strains were isolated from the central nervous system, which is an inositol rich environment. Because most of the *ITR4* gene variants are proximal to the coding region, these alterations may change expression of the *ITR4* gene, or transcript/protein stability *in vivo*, rather than abolish gene expression as occurs in the *itr4*Δ mutant. This could explain the difference between *in vitro* inositol phenotypes we observed between our clinical isolates and the mutant. It is also possible that the genetic background of the clinical isolates influences the function of the different *ITR4* gene variants, as these genes are known to be part of larger inositol acquisition and utilization pathways.

There was no clear relationship between genes that were identified in both of our statistical analyses and the gene deletion virulence in mice (five genes were significant in both, two with a significant gene deletion virulence effect, Table S8). We note, however, that although there is a good link between strain survival in mice and human virulence [24], there are two major limitations with interpretation and extrapolation of the virulence tests we performed in this study. The first is that the phenotype of a gene knockout does not necessarily recapitulate the effect of a natural point or indel mutation (e.g., [51–53]). Importantly, variants located upstream of a gene were extremely prevalent in our dataset, suggesting that they would not be phenocopied with a gene deletion if an increase in expression is required to influence the trait. The second reason for pause is that the gene knockout collection is in the KN99*α* genetic background. It has previously been shown that although ST93 and KN99*α* are both VNI strains, they are phylogenetically quite distantly related [10]. We see this distance in our own dataset: 2941 variants were present in the closely related ST93 genomes we sequenced and over 40 000 variants were present across all the genomes compared to the H99 reference strain. Genetic background is known to play a significant role on the effect of a mutation. A large study in *Saccharomyces cerevisiae* recently found that 16-42% of deletion phenotypes changed between pairs of strains, depending on the environment [54]. To fully probe the influence of the variants and genes we identified in our screen these variants need to be studied in the ST93 background. Given these limitations, we anticipate additional studies will uncover more genes from our study with an impact on pathogenesis. It would also of course be of general interest to reconstruct a knockout collection in a strain background more representative of typical clinical strains [14,28].

We purposefully chose to focus our study on strains from ST93, which was the most prevalent ST group from strains we sampled from participants in the COAT trial (~63% of all strains). ST did not significantly influence mortality (ST93: 22 patients died, 24 survived; non-ST93: 9 patients died, 16 survived; fisher-exact test *p* = 0.45). ST93 was similarly the most prevalent among advanced HIV patients in Brazil [25]. By contrast, ST93 isolates were less common than ST5 isolates among immunocompetent patients in Vietnam, and non-ST5 strains were associated with decreased mortality compared to ST5 [55]. Other studies have found no ST93 isolates [56,57]. This picture of geography having a major impact on which group is most prevalent begs the question of whether it is merely chance or the effect of selection that sorts lineages geographically.

As additional ‘genome enabled’ clinical datasets are constructed, we can hope to gain a clearer global picture about the link between broad and narrow genomic variability on clinical outcome. Our narrow analysis in the ST93 strains was possible because of the large number of patients infected with this sequence type in Uganda. Only as similar studies are performed in patient populations throughout the world, with other dominant STs, or in the context of increased genetic diversity, will we be able to determine how broadly applicable our study is to the global population of *C. neoformans*.

Statistical association techniques using human clinical data, such as those employed here, offer a complementary approach to genetic screens of mutant collections. They offer the benefit of not having to choose a particular strain background to focus your efforts (typically the reference strain), nor make decisions about which genes are likely to be the most important. For example, the genes chosen for the initial *C. neoformans* knockout collection were biased not only against genes with homologs in *S. cerevisiae*, but also against *C. neoformans-specific* genes [7]. There are also inherent biases to forward genetics methods. Here we only have the statistical power to find association with common variants. The majority of variants we sampled among our strains were singleton variants in only a single genome (Figure 1A), and some of these may well have an extremely important influence on virulence that remains undetected in our current analysis. Hence we have treated our pathogen GWAS analysis like a genetic screen, and the true utility of this type of analysis is not seen in just one study in isolation of others. The power lies in the opportunity to compare among studies of different types to find candidate genes or alleles to focus our attention on.

Our analysis did not identify variants in many of the genes that were previously identified through *in vitro* and in animal mutant screens as virulence factors in *C. neoformans*, such as genes involved in capsule formation and melanin synthesis. There could be several reasons for this result. Importantly, all of the ST93 strains analyzed were isolated from patients with cryptococcal meningitis, thus all these strains by definition are capable of causing disease and in our study the readout was not presence or absence of disease but rather the severity of disease. Previous studies may have identified virulence factors involved in the early stages of infection that impact the ability of *C. neoformans* to infect and then survive within the host, whereas our study identified virulence factors that promote or inhibit the progression of disease. Second, our analysis utilized human clinical data for association with genetic differences between strains whereas previous studies utilized surrogates, either *in vitro* conditions or animal models. By studying genetic differences in the context of human infection, we have the potential to not only define genes that promote disease in humans but also the potential to define aspects of the host-pathogen interaction that are specific to *C. neoformans* and the human host.

## Methods

### Ethics Statement

Animal experiments were done in accordance with the Animal Welfare Act, United States federal law, and NIH guidelines. Mice were handled in accordance with guidelines defined by the University of Minnesota Animal Care and Use Committee (IACUC) under protocol 1607-34001A. Participant data were collected as part of the COAT trial (clinicaltrials.gov:NCT01075152) [30,40]. All participants were enrolled in Uganda at Mulago Hospital, Makerere University, in Kampala. Written informed consent was obtained from all subjects or their proxy, and all data were de-identified. Institutional Review Board (IRB) approvals were obtained both at the University of Minnesota (0810M49622) and Makerere University.

### Strain selection

We utilized *C. neoformans* isolates collected in Uganda as part of the Cryptococcal Optimal ART Timing (COAT) trial [30]. We focused primarily on 38 UgCl (“**Ug**andan **Cl**inical”) COAT strains that had previously been MLST genotyped as sequence type 93 (ST93), the most prevalent ST group in this collection of strains [31]. An additional 18 strains from ten MLST groups were also whole genome sequenced to represent strain diversity in Ugandan clinical isolates [9].

Clinical isolates were colony purified from the CSF of participants that presented at the clinic with their first episode of cryptococcal meningitis. The ST93 clinical isolate strains were purposefully chosen to represent strains from both participants who survived (n=21) and died (n = 17). As with the parent COAT trial, survival was decreased with early ART initiation, all putative ST93 clinical isolates used for these studies were from the standard-of-care (deferred ART treatment) arm of the clinical trial. Patient infection phenotypes (i.e., clinical and cytokine parameters, Table 1) were measured on the day patients were diagnosed with cryptococcal meningitis, prior to antifungal or ART treatment. Cytokine data was log2 transformed prior to analysis, as described previously [32].

### Library Preparation and Illumina Sequencing

DNA was extracted using the CTAB DNA isolation method. Colony-purified cultures, maintained as glycerol stocks at −80°C, were inoculated into 250mL yeast peptone dextrose agar (YPD) in erlenmeyer flasks and grown overnight at 30°C with continuous shaking prior to DNA isolation.

Strains were whole-genome sequenced in two sets. In the first set, genomic DNA libraries from 16 strains were prepared by the Mayo Bioinformatics Core for 101bp paired-end sequencing. The samples were combined into two pools (Pool A: UgCl001, UgCl018, UgCl021, UgCl029, UgCl030, UgCl037, UgCl040, UgCl045, UgCl057, UgCl074, UgCl076, UgCl107; Pool B: UgCl008, UgCl032, UgCl047, UgCl065, UgCl087, UgCl093). Each pool was sequenced on a single lane of an Illumina HiSeq 20009.

In the second set, genomic DNA libraries from the 40 strains were prepared by the University of Minnesota Genomics Center for 300bp paired-end sequencing with the Illumina TruSeq DNA LT kit. The samples were combined into four pools; each pool was sequenced in a single lane of an Illumina MiSeq (Pool1: UgCl212, UgCl230, UgCl236, UgCl243, UgCl247, UgCl250, UgCl389, UgCl541, UgCl547, UgCl549; Pool2: UgCl252, UgCl255, UgCl262, UgCl291, UgCl292, UgCl300, UgCl326, UgCl332, UgCl357, UgCl360; Pool3: UgCl362, UgCl377, UgCl379, UgCl382, UgCl390, UgCl393, UgCl395, UgCl422, UgCl438, UgCl443; Pool4: UgCl447, UgCl450, UgCl461, UgCl462, UgCl466, UgCl468, UgCl495, UgCl534, UgCl535, UgCl538, UgCl546). In the second set of sequencing, the runs generated 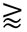 22 million pass filter reads for pools 1 and 2 and 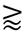 17 million pass filter reads for pools 3 and 4. In all runs >70% bases were above Q30. The average library insert size was 400-500bp.

### Variant calling

Variant calling for each strain was adapted from the Genome Analysis Toolkit (GATK v3.3.0) best practices [33–35]. For each strain the two paired-end fastq files were trimmed using trimmomatic [36] and aligned to the *C. neoformans* H99 reference genome downloaded from FungiDB (http://fungidb.org/fungidb/) on February 1, 2016 (“FungiDB-26_Cneoformans_H99_Genome.fasta”) with bwa mem [37]. The output (.SAM) files from all other strains were converted to .BAM files and sorted, duplicates were marked and indexed and a final index was built with picard tools (http://broadinstitute.github.io/picard). Variants were called for each sample with GATK HaplotypeCaller run in GVCF mode for each strain (with flags --genotyping_mode DISCOVERY --emitRefConfidence GVCF -variant_index_type LINEAR -variant_index_parameter 128000 -ploidy 1) to obtain gVCF files. GATK GenotypeGVCFs was then run to merge the 41 gVCF records. Variants were annotated with SnpEff [38] followed by GATK VariantAnnotator. SNPs and INDELs were separated into two tables from the single merged and annotated VCF file using GATK SelectVariants, VariantFiltration and VariantsToTable. Coverage across chromosomes was determined using GATK DepthOfCoverage on the sorted BAM files.

### Phylogenetic tree building

SNPhylo [39], a pipeline designed to construct phylogenetic trees from SNP data, was used to generate a PHYLIP file from the original VCF. SNPhylo reduces redundant SNP information due to linkage disequilibrium. As we knew *a priori* that our ST93 samples are highly related, we ran SNPhylo with the linkage disequilibrium flag set very high (0.99), which still reduced the number of SNPs by ~94% on each chromosome. 7,383 markers were selected in total. In SNPhylo, MUSCLE was used to perform multiple alignment and generate the PHYLIP file.

Bootstrap analysis was conducted using RAxML. 20 maximum likelihood trees were generated (-m ASC_GTRGAMMA --asc-corr=lewis) and support values from 100 bootstrap replicates were determined for the best fit ML tree (-m ASC_GTRGAMMA --asc-corr=lewis -p 3 -b 12345 -#100). Bipartitions were then drawn on the best tree (-m ASC_GTRGAMMA --asc-corr=lewis -p 3 -f b). This tree was read into R using the read.raxml command in the treeio library. Further tree visualisations were created using ggtree.

### Clinical data

Collection of clinical and immunological data were as described previously [30,40]. Clinical and immunological data used in this study are listed in Table 1. Briefly, clinical parameters of disease were participant mortality due to cryptococcosis (days post initial diagnosis), CD4+ T-cell count, cerebral spinal fluid (CSF) white blood cell count (WBC), serum and CSF protein levels, HIV viral load, CSF *Cryptococcus* clearance rate of early fungicidal activity (EFA), and lateral flow assay (LFA) measurement of cryptococcal antigen titer (Immy Inc., Norman, Oklahoma). As immunological data, CSF levels of 19 cytokines and chemokines (granulocyte colony-stimulating factor [G-CSF], granulocyte macrophage colony-stimulating factor [GM-CSF], interferon-γ, tumor necrosis factor [TNF]—α, interleukin [IL]–1β, IL-2, IL-4, IL-5, IL-6, IL-7, IL-8, IL-10, IL-12, IL-13, IL-17, MCP-1 [CCL2], macrophage inflammatory protein [MIP]–1α [CCL3], MIP-1β [CCL4], VEGF) were analyzed. We refer to these cytokines and chemokines collectively as “cytokines”.

*In vitro* assays were also performed on the clinical isolates. Drug resistance assays for fluconazole and amphotericin B were as described previously [31,41]. MH-S macrophage cell cultures were used to determine *C. neoformans* cell uptake by macrophages. Briefly, 5×10^5^ MH-S cells per well were incubated at 37°C with 5% CO_2_ for 2 hours in a 96-well culture plate to allow adherence. *C. neoformans* cultures were grown overnight in Dulbecco’s modified eagle medium (DMEM) supplemented with 2% glucose, collected by centrifugation, washed, and resuspended in 0.1% Uvitex solution for 10 minutes. Cells were then collected by centrifugation, washed, and 5×10^5^ cells and 4*μ*g E1 anti-GXM antibody [42] were added to each well in the MH-S culture plate. After two hours of co-incubation, the culture plate was centrifuged to collect cells, spent media was decanted and cells were washed to remove extracellular *C. neoformans* cells. Samples were then resuspended in 0.25% Trypsin in EDTA for 15 minutes to release the adherent cells from the wells, fixed with 3.7% formaldehyde for 30 minutes on ice. Samples were then stained with a second anti-GXM antibody (m18b7) conjugated to the AlexaFluor 488 fluorophore (1:2000) and PE-labelled CD45 (1:100) in PBS with 1 *μ*g/ml bovine serum albumin (BSA and 2 mM Tris-HCl. Cells were analyzed on a BD LSRII flow cytometer (BD Biosciences, Inc) and data were analyzed using FlowJo software. Gating on Uvitex, CD45, and m18b7 allowed differentiation free *C. neoformans* cells (Uvitex+, CD45-), free macrophages (Uvitex-, CD45+), macrophages with intracellular *C. neoformans* (Uvitex+. CD45+, m18b7-), and macrophages with extracellular *C. neoformans* (Uvitex+, CD45+, m18b7+). To analyze cell wall chitin content, *C. neoformans* cells were grown in DMEM supplemented with 2% glucose, 10% FBS, 1% Pen-Strep, and beta-mercaptoethanol (1ml/1L) at 37°C overnight, and then fixed for 30 minutes in 3.7% formaldehyde. Cell concentration was adjusted to 1×10^6^ cells/ml, stained with 1*μ*g/ml calcofluor white (Sigma Aldrich) in PBS for 5 minutes at 25°C, then wash with PBS. Median Calcofluor white fluorescence intensity was then determined for each strain by flow cytometric analysis of the cell population on an LSR II Fortessa flow cytometer.

Biomarkers analyzed as continuous variables were log2 transformed for normalization, analyzed, and then back-transformed to geometric mean values. All “mean” biomarker values are geometric means. Low (“out of range”) measurements were set to half of the manufacturer’s listed assay limit of detection (LOD). CSF biomarkers with substantial proportions (≥40%) of undetectable values at diagnosis (IL-2, IL-1β, IL-5, CCL22) were analyzed as categorical variables: “detectable” (values greater than the LOD) versus “nondetectable (values lower than the LOD). CSF white blood cell count (WBC) was analyzed as absolute values and also classified as “normal” versus “elevated” (<5 vs ≥5 cells/*μ*L, respectively).

### Survival Curves

Survival curves were performed in three experiments. Experiment one (E1) tested the virulence of KN99α with the following genes deleted: CNAG_00363, CNAG_02176, CNAG_04373, CNAG_04535, CNAG_04922, CNAG_05662, CNAG_05663, CNAG_05913, CNAG_06169, CNAG_06332, CNAG_06574, CNAG_06704, CNAG_06876, and CNAG_07837. For E1, five C57BL/6 mice per group were anesthetized by intraperitoneal pentobarbital injection and inoculated intranasally with 5 × 10^4^ cells suspended in 50 μl PBS, whereas E2 and E3 used five C57BL/6 mice per group were anesthetized and inoculated intranasally with 1 × 10^4^ cells suspended in 50 μl PBS. Animals were monitored for morbidity and sacrificed with carbon dioxide when endpoint criteria were reached. Endpoint criteria were defined as 20% total body weight loss, loss of two grams of weight in two days, or symptoms of neurological damage. On day 34, the remaining mouse was sacrificed. Lungs and brain were removed and homogenized in 4 mL or 2 mL PBS, respectively. Serial dilutions of the lungs and the entire homogenized brain were plated on YPD with chloramphenicol. CFUs were counted after 48 hours.

Significance was determined using the *survfit* command from the R package survival [43]. Kaplan-Meier estimators from each knockout strain were compared against the KN99α strain measured in the relevant experiment. P-values were obtained by comparing the two curves using the G-rho family log-rank test [44], implemented with the *survdiff* function.

### ITR4 Survival Curve

Ten C57BL/6 mice per group were anesthetized and inoculated intranasally with 1 × 10^3^ KN99α, *itr4Δ*, or *itr4Δ:ITR4* cells suspended in 50 μl PBS. Animals were treated as described above. The *itr4Δ* that survived the infection initially showed early signs of disease (minor weight loss, reduced activity) but regain weight at later timepoints. On day 44, the mice were sacrificed. Lungs and brain were collected from each mouse to determine fungal burden, and processed as described above.

For determination of CFUs at 7 days post-infection, 4 C57BL/6 mice per group were anesthetized and inoculated intranasally with 1 × 10^3^ KN99α, *itr4Δ*, or *itr4Δ:ITR4* cells suspended in 50 μl PBS. After seven days, the mice were sacrificed, and lungs and brain were collected and processed as described above.

### Inositol Growth assays

Yeast cells of *C. neoformans* wild type strain *KN99α, itr4Δ* mutant, and clinical strains were cultured in YPD medium overnight. Concentrations of overnight cultures were determined by measuring the optical density at 600 nm (OD600) and adjusted to the same cell density. Serial 10-fold dilutions were prepared, and 5 μl of each dilution was spotted on YNB plates with 1% glucose, 1% inositol, or 1% glucose and 1% inositol. Plates were then incubated at 30°C or 37°C for 48 h before photography. The assay was repeated at least three times with similar results.

### Inositol uptake assay

The inositol uptake assay was performed following the previously published method [60]. In brief, the *Cryptococcus* strains were grown in YPD liquid cultures overnight at 30°C. Cells were diluted in YPD to an OD600 of 1.0, grown at 30°C, and collected at an OD600 of 5.0 by centrifugation at 2,600 × *g* for 5 min. Cells were then washed twice with PBS at 4°C and resuspended in 2% glucose to a final concentration of 2 × 10^8^ cells/ml as determined by a hemacytometer. For the uptake assay, the reaction mixture (200 *μ*l) contained 2% glucose, 40 mM citric acid-KH_2_PO_4_ (pH 5.5), 0.15 *μ*M myo-[2-^3^H]-inositol (1 μCi/μl; MP BioMedicals). Additional 200 μM unlabeled inositol (Sigma-Aldrich) was added to the reactions for competition assays. Equal volumes of the reaction and cell mixtures (60 *μ*l each) were warmed to 30°C and mixed for the uptake assay, which was performed for 10 min at 30°C. As negative controls, mixtures were kept at 0°C (on ice) during the 10-min incubation. Aliquots of 100 *μ*l were removed and transferred onto prewetted Metricel filters (1.2 *μ*m) on a vacuum manifold. The filters were washed four times each with 2 ml of ice-cold water. The washed filters were removed and added to liquid scintillation vials for measurements on a PerkinElmer TRI-CARB 2900TR scintillation counter.

### Data Availability

All data and scripts are available at GitHub at https://github.com/acgerstein/UgClGenomics

## Supporting Information Legends

### Supplemental Tables

Table S1. Genes, hypothetical RNAs, and intergenic regions with variants that are present in all ST93 genomes

Table S2. Genes with at least 10 variants present in all ST93 genomes Table S3. ST93A and ST93B clade-specific variants

Table S4. Statistical analysis of ST93 clade-specific associations with quantitative infection phenotypes

Table S5. Phenotypes measured from patients enrolled in the COAT trial (clinical and cytokines) and *in vitro*.

Table S6. Significant variants in genes and hypothetical RNAs with quantitative infection phenotypes based on class designation

Table S7. Logistic regression analysis of all significant variants in genes and hypothetical RNAs associated with quantitative infection phenotypes

Table S8. The majority of genes associated with quantitative infection phenotypes are uncharacterized

### Supplemental Figure Legends

**Figure S1. Clade-specific differences in phenotype**. Bar indicates median value.

**Figure S2. PCA analysis**. A) Each dashed line represents one of 20 randomized trials. B) There was no association between PC1 or PC2 and clade.

**Figure S3. KN99α deletion strain virulence in mice**. Groups of five 6-8 week old C57Bl/6 mice were infected intranasally with 5 × 10^4^ cells. Progression to severe morbidity was monitored for 35 days and mice were sacrificed when endpoint criteria were reached. Strains were tested in two separate experiments, E1 or E2, respectively. The deletions strains were compared against the KN99α strain in the same experiment.

**Figure S4. Growth at 7 days post-infection**. Groups of four 6-8 week old C57Bl/6 mice were infected intranasally with 1 × 10^3^ cells. Mice were sacrificed at 7 days post infection, lungs homogenized in 4 ml of PBS, and serial dilutions plated on YPD with cholamphenicol medium. Colony forming units were enumerated at 48 hours.

## References

1. Rajasingham R, Smith RM, Park BJ, Jarvis JN, Govender NP, Chiller TM, et al. Global burden of disease of HIV-associated cryptococcal meningitis: an updated analysis. Lancet Infect Dis. 2017;17: 873–881.

2. Cowen LE, Sanglard D, Howard SJ, Rogers PD, Perlin DS. Mechanisms of Antifungal Drug Resistance. Cold Spring Harb Perspect Med. 2014;5: a019752.

3. Butts A, Krysan DJ. Antifungal drug discovery: something old and something new. PLoS Pathog. 2012;8: e1002870.

4. Azevedo RVDM, Rizzo J, Rodrigues ML. Virulence Factors as Targets for Anticryptococcal Therapy. J Fungi (Basel). 2016;2. doi:10.3390/jof2040029

5. Brunke S, Mogavero S, Kasper L, Hube B. Virulence factors in fungal pathogens of man. Curr Opin Microbiol. 2016;32: 89–95.

6. Gerstein AC, Nielsen K. It’s not all about us: evolution and maintenance of *Cryptococcus* virulence requires selection outside the human host. Yeast. 2017;34: 143–154.

7. Liu OW, Chun CD, Chow ED, Chen C, Madhani HD, Noble SM. Systematic genetic analysis of virulence in the human fungal pathogen *Cryptococcus neoformans*. Cell. 2008;135: 174–188.

8. Shea JM, Kechichian TB, Luberto C, Del Poeta M. The Cryptococcal Enzyme Inositol Phosphosphingolipid-Phospholipase C Confers Resistance to the Antifungal Effects of Macrophages and Promotes Fungal Dissemination to the Central Nervous System. Infect Immun. 2006;74: 5977–5988.

9. Wiesner DL, Moskalenko O, Corcoran JM, McDonald T, Rolfes MA, Meya DB, et al. Cryptococcal genotype influences immunologic response and human clinical outcome after meningitis. MBio. 2012;3. doi:10.1128/mBio.00196-12

10. Beale MA, Sabiiti W, Robertson EJ, Fuentes-Cabrejo KM, O’Hanlon SJ, Jarvis JN, et al. Genotypic Diversity Is Associated with Clinical Outcome and Phenotype in Cryptococcal Meningitis across Southern Africa. PLoS Negl Trop Dis. 2015;9: e0003847.

11. Tefsen B, Grijpstra J, Ordonez S, Lammers M, van Die I, de Cock Cock H. Deletion of the CAP10 gene of *Cryptococcus neoformans* results in a pleiotropic phenotype with changes in expression of virulence factors. Res Microbiol. 2014;165: 399–410.

12. Griffiths EJ, Kretschmer M, Kronstad JW. Aimless mutants of *Cryptococcus neoformans:* failure to disseminate. Fungal Biol Rev. 2012;26: 61–72.

13. Sabiiti W, Robertson E, Beale MA, Johnston SA, Brouwer AE, Loyse A, et al. Efficient phagocytosis and laccase activity affect the outcome of HIV-associated cryptococcosis. J Clin Invest. 2014;124: 2000–2008.

14. Motaung TE. *Cryptococcus neoformans* mutant screening: a genome-scale’s worth of function discovery. Fungal Biol Rev. 2018;32: 181–203.

15. Desalermos A, Tan X, Rajamuthiah R, Arvanitis M, Wang Y, Li D, et al. A multi-host approach for the systematic analysis of virulence factors in *Cryptococcus neoformans*. J Infect Dis. 2015;211: 298–305.

16. Kwon-Chung KJ, Boekhout T, Fell JW, Diaz M. (1557) Proposal to Conserve the Name *Cryptococcus gattii* against *C. hondurianus* and *C. bacillisporus* (Basidiomycota, Hymenomycetes, Tremellomycetidae). Taxon. International Association for Plant Taxonomy (IAPT); 2002;51: 804–806.

17. Meyer W, Marszewska K, Amirmostofian M, Igreja RP, Hardtke C, Methling K, et al. Molecular typing of global isolates of *Cryptococcus neoformans* var. *neoformans* by polymerase chain reaction fingerprinting and randomly amplified polymorphic DNA—a pilot study to standardize techniques on which to base a detailed epidemiological survey. Appl Theor Electrophor. Wiley Online Library; 1999;20: 1790–1799.

18. Hagen F, Lumbsch HT, Arsic Arsenijevic V, Badali H, Bertout S, Billmyre RB, et al. Importance of Resolving Fungal Nomenclature: the Case of Multiple Pathogenic Species in the *Cryptococcus* Genus. mSphere. 2017;2. doi:10.1128/mSphere.00238-17

19. Hagen F, Khayhan K, Theelen B, Kolecka A, Polacheck I, Sionov E, et al. Recognition of seven species in the *Cryptococcus gattii/Cryptococcus neoformans* species complex. Fungal Genet Biol. 2015;78: 16–48.

20. Meyer W, Firacative C, Trilles L, Ferreira-Paim K, ISHAM Working Group for Genotyping of C. neoformans and C. gattii, abstr. MLST database for *Cryptococcus neoformans* and *C. gattii*. 19th ISHAM Congress. Melbourne, Australia, 4 to 8 May 2015.

21. Kwon-Chung KJ, Bennett JE, Wickes BL, Meyer W, Cuomo CA, Wollenburg KR, et al. The Case for Adopting the “Species Complex” Nomenclature for the Etiologic Agents of Cryptococcosis. mSphere. 2017;2. doi:10.1128/mSphere.00357-16

22. Desnos-Ollivier M, Patel S, Raoux-Barbot D, Heitman J, Dromer F, French Cryptococcosis Study Group. Cryptococcosis Serotypes Impact Outcome and Provide Evidence of *Cryptococcus neoformans* Speciation. MBio. 2015;6: e00311.

23. Dixit A, Carroll SF, Qureshi ST. Cryptococcus gattii: An Emerging Cause of Fungal Disease in North America. Interdiscip Perspect Infect Dis. 2009;2009: 840452.

24. Mukaremera L, MacDonald TR, Nielsen JN, Molenaar C, Akampulira A, Schutz C, Taseera K, Muzoora C, Meintjes G, Meya DB, Boulware DR, and Nielsen K. The mouse inhalation model of *Cryptococcus neoformans* infection recapitulates strain virulence in humans and shows closely related strains can possess differential virulence. Infect Immun. 2019;in press.

25. Andrade-Silva LE, Ferreira-Paim K, Ferreira TB, Vilas-Boas A, Mora DJ, Manzato VM, et al. Genotypic analysis of clinical and environmental *Cryptococcus neoformans* isolates from Brazil reveals the presence of VNB isolates and a correlation with biological factors. PLoS One. 2018;13: e0193237.

26. Ferreira-Paim K, Andrade-Silva L, Fonseca FM, Ferreira TB, Mora DJ, Andrade-Silva J, et al. MLST-based population genetic analysis in a global context reveals clonality amongst *Cryptococcus neoformans* var. *grubii* VNI isolates from HIV patients in Southeastern Brazil. PLoS Negl Trop Dis. Public Library of Science; 2017;11: e0005223.

27. Khayhan K, Hagen F, Pan W, Simwami S, Fisher MC, Wahyuningsih R, et al. Geographically structured populations of *Cryptococcus neoformans* Variety *grubii* in Asia correlate with HIV status and show a clonal population structure. PLoS One. 2013;8: e72222.

28. Ashton PM, Thanh LT, Trieu PH, Van Anh D, Trinh NM, Beardsley J, et al. Genomics of *Cryptococcus neoformans* [Internet]. bioRxiv. 2018. p. 356816. doi:10.1101/356816

29. Desjardins CA, Giamberardino C, Sykes SM, Yu C-H, Tenor JL, Chen Y, et al. Population genomics and the evolution of virulence in the fungal pathogen *Cryptococcus neoformans*. Genome Res. 2017;27: 1207–1219.

30. Boulware DR, Meya DB, Muzoora C, Rolfes MA, Huppler Hullsiek K, Musubire A, et al. Timing of antiretroviral therapy after diagnosis of cryptococcal meningitis. N Engl J Med. 2014;370: 2487–2498.

31. Smith KD, Achan B, Hullsiek KH, McDonald TR, Okagaki LH, Alhadab AA, et al. Increased Antifungal Drug Resistance in Clinical Isolates of *Cryptococcus neoformans* in Uganda. Antimicrob Agents Chemother. 2015;59: 7197–7204.

32. Boulware DR, Meya DB, Bergemann TL, Wiesner DL, Rhein J, Musubire A, et al. Clinical features and serum biomarkers in HIV immune reconstitution inflammatory syndrome after cryptococcal meningitis: a prospective cohort study. PLoS Med. 2010;7: e1000384.

33. McKenna A, Hanna M, Banks E, Sivachenko A, Cibulskis K, Kernytsky A, et al. The Genome Analysis Toolkit: a MapReduce framework for analyzing next-generation DNA sequencing data. Genome Res. 2010;20: 1297–1303.

34. DePristo MA, Banks E, Poplin R, Garimella KV, Maguire JR, Hartl C, et al. A framework for variation discovery and genotyping using next-generation DNA sequencing data. Nat Genet. 2011;43: 491–498.

35. Bateman A, Pearson WR, Stein LD, Stormo GD, Yates JR III, editors. From FastQ Data to High-Confidence Variant Calls: The Genome Analysis Toolkit Best Practices Pipeline: The Genome Analysis Toolkit Best Practices Pipeline. Current Protocols in Bioinformatics. Hoboken, NJ, USA: John Wiley & Sons, Inc.; 2002. pp. 11.10.1–11.10.33.

36. Bolger AM, Lohse M, Usadel B. Trimmomatic: a flexible trimmer for Illumina sequence data. Bioinformatics. 2014;30: 2114–2120.

37. Li H, Durbin R. Fast and accurate long-read alignment with Burrows-Wheeler transform. Bioinformatics. 2010;26: 589–595.

38. Cingolani P, Platts A, Wang LL, Coon M, Nguyen T, Wang L, et al. A program for annotating and predicting the effects of single nucleotide polymorphisms, SnpEff: SNPs in the genome of Drosophila melanogaster strain w1118; iso-2; iso-3. Fly. 2012;6: 80–92.

39. Lee T-H, Guo H, Wang X, Kim C, Paterson AH. SNPhylo: a pipeline to construct a phylogenetic tree from huge SNP data. BMC Genomics. 2014;15: 162.

40. Scriven JE, Rhein J, Hullsiek KH, von Hohenberg M, Linder G, Rolfes MA, et al. Early ART After Cryptococcal Meningitis Is Associated With Cerebrospinal Fluid Pleocytosis and Macrophage Activation in a Multisite Randomized Trial. J Infect Dis. 2015;212: 769–778.

41. Nielsen K, Vedula P, Smith KD, Meya DB, Garvey EP, Hoekstra WJ, et al. Activity of VT- 1129 against *Cryptococcus neoformans* clinical isolates with high fluconazole MICs. Med Mycol. 2017;55: 453–456.

42. Dromer F, Salamero J, Contrepois A, Carbon C, Yeni P. Production, characterization, and antibody specificity of a mouse monoclonal antibody reactive with *Cryptococcus neoformans* capsular polysaccharide. Infect Immun. 1987;55: 742–748.

43. Therneau TM. A Package for Survival Analysis in S [Internet]. 2015. Available: https://CRAN.R-project.org/package=survival

44. Harrington DP, Fleming TR. A Class of Rank Test Procedures for Censored Survival Data. Biometrika. [Oxford University Press, Biometrika Trust]; 1982;69: 553–566.

45. Ashton APM, Thanh LT, Trieu PH, Van Anh D, Trinh NM, Beardsley J. 1 Genomics of *Cryptococcus neoformans*. doi:10.1101/356816

46. Hirschhorn JN, Daly MJ. Genome-wide association studies for common diseases and complex traits. Nat Rev Genet. 2005;6: 95–108.

47. Little RJ, D’Agostino R, Cohen ML, Dickersin K, Emerson SS, Farrar JT, et al. The prevention and treatment of missing data in clinical trials. N Engl J Med. 2012;367: 1355–1360.

48. Chun CD, Madhani HD. Chapter 33 - Applying Genetics and Molecular Biology to the Study of the Human Pathogen Cryptococcus neoformans. Elsevier Inc; 2010.

49. Luberto C, Martinez-Mariño B, Taraskiewicz D, Bolaños B, Chitano P, Toffaletti DL, et al. Identification of App1 as a regulator of phagocytosis and virulence of *Cryptococcus neoformans*. J Clin Invest. 2003;112: 1080–1094.

50. Derengowski L da S, Paes HC, Albuquerque P, Tavares AHFP, Fernandes L, Silva-Pereira I, et al. The transcriptional response of Cryptococcus neoformans to ingestion by *Acanthamoeba castellanii* and macrophages provides insights into the evolutionary adaptation to the mammalian host. Eukaryot Cell. 2013;12: 761–774.

51. Sareila O, Hagert C, Rantakari P, Poutanen M, Holmdahl R. Direct Comparison of a Natural Loss-Of-Function Single Nucleotide Polymorphism with a Targeted Deletion in the Ncf1 Gene Reveals Different Phenotypes. PLoS One. 2015;10: e0141974.

52. Housden BE, Muhar M, Gemberling M, Gersbach CA, Stainier DYR, Seydoux G, et al. Loss-of-function genetic tools for animal models: cross-species and cross-platform differences. Nat Rev Genet. 2017;18: 24–40.

53. Tamae C, Liu A, Kim K, Sitz D, Hong J, Becket E, et al. Determination of antibiotic hypersensitivity among 4,000 single-gene-knockout mutants of *Escherichia coli*. J Bacteriol. 2008;190: 5981–5988.

54. Galardini M, Busby BP, Vieitez C, Dunham AS, Typas A, Beltrao P. The impact of the genetic background on gene deletion phenotypes in *Saccharomyces cerevisiae* [Internet]. bioRxiv. 2018. p. 487439. doi:10.1101/487439

55. Day JN, Qihui S, Thanh LT, Trieu PH, Van AD, Thu NH, et al. Comparative genomics of *Cryptococcus neoformans* var. *grubii* associated with meningitis in HIV infected and uninfected patients in Vietnam. PLoS Negl Trop Dis. 2017;11: e0005628.

56. Fernandes KE, Brockway A, Haverkamp M, Cuomo CA, van Ogtrop F, Perfect JR, et al. Phenotypic Variability Correlates with Clinical Outcome in Cryptococcus Isolates Obtained from Botswanan HIV/AIDS Patients. MBio. 2018;9. doi:10.1128/mBio.02016-18

57. Miglia KJ, Govender NP, Rossouw J, Meiring S, Mitchell TG, Group for Enteric, Respiratory and Meningeal Disease Surveillance in South Africa. Analyses of pediatric isolates of *Cryptococcus neoformans* from South Africa. J Clin Microbiol. 2011;49: 307–314.

58. Ereshefsky M. Microbiology and the Species Problem. Biology and Philosophy. 2010; 25: 67–79.

59. Xue C, Liu L, Li W, Liu I, Kronstad JW, Seyfeng A, Heitman. Role of an expanded inositol transporter repertoire in Cryptococcus neoformans sexual reproduction and virulence. mBio. 2010; 1: e00084–10.

60. Wang Y, Liu T, Delmas G, Park S, Perlin D, Xue C. Two major inositol transporters and their role in cryptococcal virulence. Euk. Cell. 2011; 10: 618–628.

